# A cell and transcriptome atlas of human arterial vasculature

**DOI:** 10.1101/2024.09.10.612293

**Authors:** Quanyi Zhao, Albert Pedroza, Disha Sharma, Wenduo Gu, Alex Dalal, Chad Weldy, William Jackson, Daniel Yuhang Li, Yana Ryan, Trieu Nguyen, Rohan Shad, Brian T. Palmisano, João P. Monteiro, Matthew Worssam, Alexa Berezwitz, Meghana Iyer, Huitong Shi, Ramendra Kundu, Lasemahang Limbu, Juyong Brian Kim, Anshul Kundaje, Michael Fischbein, Robert Wirka, Thomas Quertermous, Paul Cheng

## Abstract

Contiguous arterial segments show different propensities for different vascular pathologies, yet mechanisms explaining these fundamental differences remain unknown. We sought to build a transcriptomic, cellular, and spatial atlas of human arterial cells across multiple different arterial segments to understand these underlying differences.

Analysis of multiple isogenic arterial segments from healthy donors reveals a significant stereotyped pattern of cell type-specific segmental heterogeneity in healthy arteries. Combining single cell analysis with spatial transcriptomic data reveals cellular heterogeneity not captured by commonly used cell-type marker genes. Determinants of arterial transcriptomic identities are predominantly encoded in fibroblasts and smooth muscle cells (SMC), and their differentially expressed genes are particularly enriched for different vascular disease-associated genetic risk-loci and risk-genes. Adventitial fibroblast-specific heterogeneity in gene expression coincides with a disproportionally large number of vascular disease genetic signals, suggesting a previously unrecognized role for this cell type in disease risk. Adult arterial cells from different segments cluster not by anatomical proximity, but by embryonic origin. Global regulon analysis of disease related segment-specific gene expression program in fibroblast and SMC enriches for binding sites of transcription factors that are developmental master regulators whose expression persists into adulthood, suggesting an important functional role of the same developmental master regulators in adult gene expression and disease. Lastly, non-coding transcriptomes across arterial cells contain extensive variation in lncRNAs expressed in cell type- and segment-specific patterns, rivaling heterogeneity in protein coding transcriptomes. Differentially expressed LncRNA demonstrate enrichment for non-coding genetic signals for vascular diseases, suggesting a potential global role of segmental specific LncRNAs in regulating inherited human vascular disease risk.

## Introduction

Human vascular diseases are the leading cause of morbidity and mortality worldwide^1^. Different segments of the human aorta and its branches have seemingly identical morphology and structures, yet have dramatically distinct propensity for different vascular diseases^2–4^. Atherosclerotic disease frequently influences coronaries and carotids, while relatively sparing the descending thoracic aorta^5,6^. Hereditary aneurysmal diseases frequently affect the aortic root and ascending aorta but not other aortic segments^7,8^. Environmental triggers such as tobacco smoking and methamphetamine abuse disproportionally bring on pathologies in the infra-renal segments of the aorta and pulmonary arteries respectively^9,10^. Even inflammatory or infectious aortopathies, such as Takayasu’s vasculitis, Kawasaki’s diseases, or syphilis aortitis, have well documented segmental preferences^11–13^. While the differences in disease propensity are well documented, the underlying causal mechanisms are less clear. Prevalent hypotheses include differential exposure to life-long hemodynamic forces, along with distinct cellular lineages from different embryonic origins^14,15^. Understanding the cellular and molecular basis for the differences between these arteries may open the door to therapies for a variety of vascular diseases.

Lineage tracing data primarily from several animal models suggest a distinct pattern of embryonic origins for the different arteries that appears conserved through vertebrates^15–18^. The coronary artery is thought to derive from several sources^19,20^. The endothelium from coronary arteries can be traced to two distinct origins, the sinus venosus and the endocardium. Coronary endothelium is thought to instruct the surrounding mesenchyme, which consists of cells originating from the proepicardial organ, to turn into smooth muscle cells. ^21,22^. The ascending aorta is thought to be made primarily of at least two populations of embryonic progenitor cells, arising from both the second heart field as well as the neural crest^21,23^. The two populations of cells mix in a distinct pattern, ending before the arch, where most smooth muscle cells are thought to come from neural crest cells of the 4th pharyngeal arch. The carotid artery arises from the 3rd pharyngeal arch. The pulmonary artery (PA) arises from the 6th pharyngeal arch, the thoracic descending aorta is thought to arise from the somitic mesoderm, and splenic mesoderm thought to be responsible for the iliac and femoral arteries. These distinct developmental origins have long been postulated to be causal for risk related to various diseases, as different segments have been shown to have differential expression of selected genes as studied with bulk RNA isolation^24,25^. Transplantation experiments in mammals have suggested arterial segments retain the disease propensities of their original site, suggesting intrinsic behavior of segments accounts for their disease propensity^26,27^. However, whether the same embryological developmental pattern exists in humans is unknown. In addition, the extent to which developmental origin influences cellular gene expression and behavior of adult vessels remains elusive.

To gain insight into these fundamental observations on differential arterial disease risk, we generated a human arterial cellular atlas spanning multiple segments of the arterial circulation. Single cell transcriptomic analysis revealed that different arterial segments exhibit distinct gene expression patterns in the adult. The distinct transcriptomic signatures were most discrepant between fibroblast and smooth muscle cells (SMC) from different vascular beds, while gene expression was relatively similar between endothelial cells and residential inflammatory cells between the vascular beds. Taken together, these data suggest that vascular identity is encoded mostly in the vascular fibroblast and smooth muscle cell populations.

## Results

To generate a baseline human vascular cellular atlas simultaneously from multiple arterial sites from the same person, we collaborated with the organ donor network to collect arterial tissues from organ donors. Up to 9 distinct arterial sites (ascending thoracic aorta (ASC), aortic arch (ARCH), descending thoracic aorta (DESC), infra-renal abdominal aorta (INFRARENAL), iliac artery (ILIAC), coronary artery (RCA), carotid artery (CAROTID), pulmonary artery (PA), and aortic root (ROOT)) were chosen to be evaluated due to their different embryonic origins and disease propensities. Available segments were collected using identical surgical techniques by a dedicated team of cardiothoracic surgeons to minimalize surgical variability (Figure 1A) and processed in an identical manner from 6 different patients. Tissues were dissociated into single cells using an enzymatic digestion approach we developed to overcome the dense vascular connective tissue (See STAR Methods) ^28–30^. This method provided for vascular smooth muscle cells to be predominantly captured, consistent with histological cellular composition ratios, while single-cell RNA quality and library complexity were well preserved. Not all sites were collected from each patient as certain segments were retained along with organs that were utilized for patient care. A total of 187,958 cells were obtained. Gene expression was normalized to account for sequencing depth and mitochondrial/ribosomal content, enabling cross-sample comparison. The data were visualized in two dimensions using UMAP, revealing distinct cell populations and transcriptional heterogeneity.

**Figure 1.**
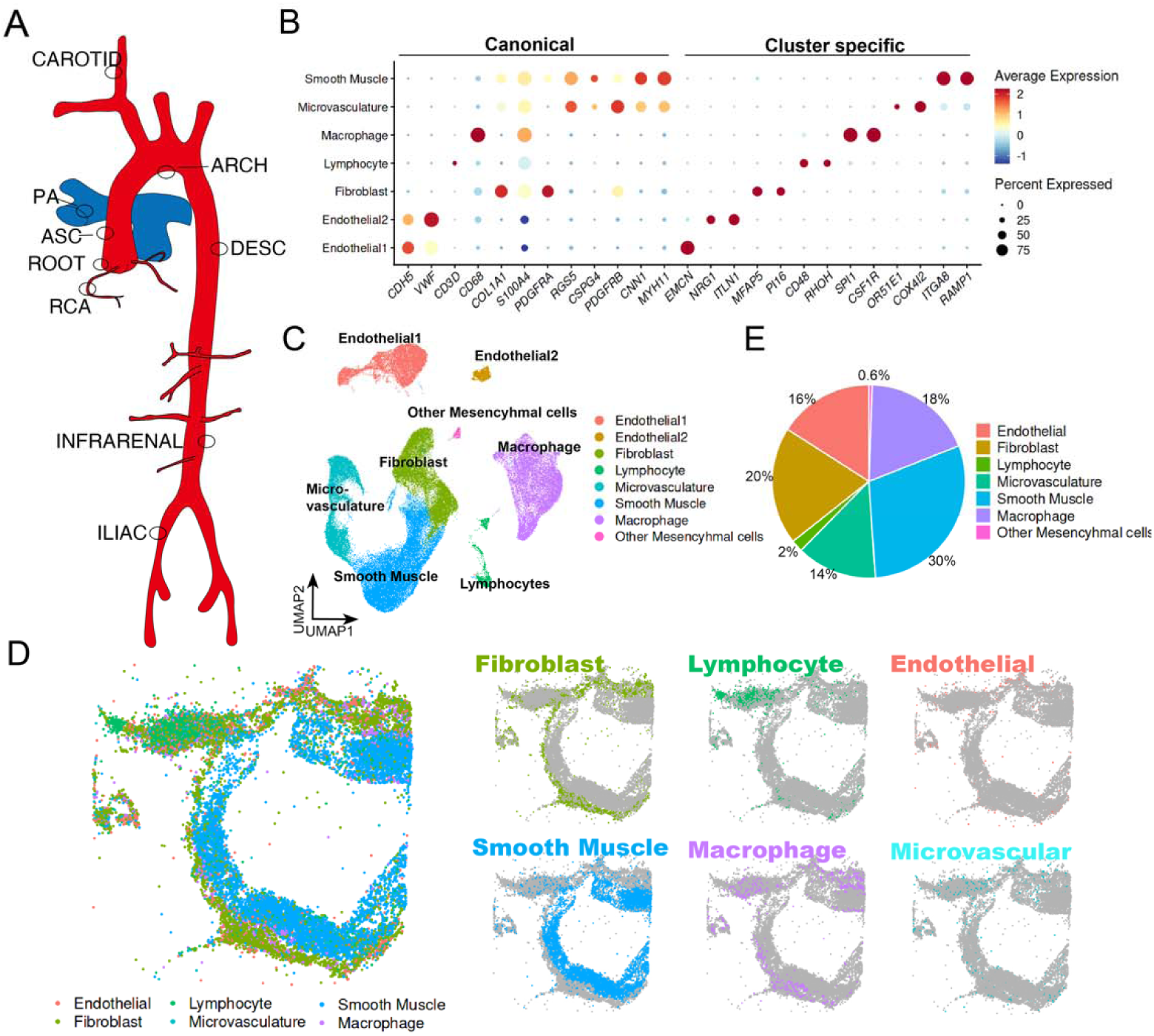
Single cell RNA and spatial landscape of human arteries. (A) Schematic illustration depicting the strategy for sampling eight anatomically distinct human arterial sites, to capture the heterogeneity of vascular cell populations across the arterial tree. (B) Heatmap showing the expression patterns of canonical markers and cluster-specific genes across all identified cell populations. Representative markers for major vascular and immune cell types are included to validate the cell type annotations. (C) Uniform Manifold Approximation and Projection (UMAP) plot of single-cell RNA-sequencing data, demonstrating distinct clustering of cells according to transcriptional profiles. Clusters are annotated by inferred cell types, including endothelial cells, smooth muscle cells, fibroblasts, immune cells, and others. (D) Spatial transcriptomics analysis of the right coronary artery (RCA). The left panel shows spatially resolved cell type annotations overlaid on tissue morphology, while the right panel presents a split-channel view, highlighting the distribution of individual cell types within the same tissue section. (E) Pie chart summarizing the overall composition of cell types in the entire single-cell RNA-seq dataset, indicating relative abundance and diversity of arterial cell populations. Abbreviations: ascending thoracic aorta (ASC), aortic arch (ARCH), descending thoracic aorta (DESC), infra-renal abdominal aorta (INFRARENAL), iliac artery (ILIAC), coronary artery (RCA), carotid artery (CAROTID), pulmonary artery (PA), and aortic root (ROOT) See also Figure S1.

The cell type identities were then identified first via unsupervised low dimensional clustering of the transcriptomic data. To avoid over-clustering, we initiated the analysis using very low-resolution clustering, which enables the identification of known distinct cell types, such as endothelial cells and smooth muscle cells. Visualization of the combined scRNA data revealed the presence of all the expected major cell types in the human arteries, namely endothelial cells (EC), smooth muscle cells (SMC), fibroblasts, microvascular cells and immune cells. These cell types are represented using existing canonical markers of each population (Figures 1B and S1A). The largest component of the variance in the dataset came from markers of different cell types, though the data contained a small level of expected inter-person variation such as gender-specific differences (Figure S1B). The resultant data are presented here (Figures 1C and S1C), and will be used as the main dimensional-reduction framework for data visualization.

### Cells of the Human Arteries

We proceeded to identify each of these specific populations of cells found in our scRNAseq data. Currently, the cellular populations in the vessels are often defined by positive staining of one or two marker genes (i.e. *CDH5* for endothelial cells), which relies on the intrinsic assumption of “marker genes” being specific, and can fail to capture unexpected heterogeneity^17,32–35^. This had led to significant debate regarding the presence and identity of certain cell types, such as vascular mesenchymal stem cells, endothelial progenitor cells, adventitial progenitor cells, pericytes, fibrocytes, fibromyocytes, modulated smooth muscle cells, fibroblasts etc^17,28,32–36^. Discrete populations of cells should have distinct transcriptomic signatures that can be identified a priori utilizing only gene expression data. Thus, we started with a very conservative clustering resolution to identify principal cell types. We then identified cluster specific gene markers, and contrasted them with the canonical “cell type markers” classically utilized to identify SMC, EC, immune cells, fibroblast, pericytes, and various progenitor cells (Figure 1B) to contrast the specificity of historical and empirically determined cell type markers. This preliminary analysis demonstrated high fidelity of canonical endothelial (*CDH5, VWF*) and immune cell markers (*CD3D, CD68*), but poor specificity of several frequently utilized “pericyte” (*RGS5, CSPG4, PDGFRB*) and “fibroblast” markers (*COL1A1, FSP1/S100A4, PDGFRA*), with robust expression seen in multiple populations (Figure S1A). To further confirm the fidelity of different cell clusters, we then mapped each clusters spatially with Slideseq (Figures 1D and S1D). Each of these distinct clusters have spatially distinct locations, with fibroblast being found mostly in the adventitia, whereas two clusters expressing canonical SMC markers (MYH11) are naturally divided into two populations that spans intima/media (termed “Smooth muscle cells’ in figure) and adventitia (termed “microvascular cells” in figure). Each of the clusters are represented in all the donor and site samples (Figures S1E and S1F). The relative proportion of each population of cells across the different arterial segments appears to be mostly constant, with SMCs making up (30%), Fibroblasts (20%), Endo (16%), Microvascular (14%), Macrophage (18%) (Figure 1E; Table S1). No sites had statistically significantly more of one cell type compared to another.

### Smooth Muscle Cells of Great Arteries

Histologically, the largest population of cells in the great arteries are SMCs. These cells make up the majority of the arterial wall^28,36,37^, and accounts for ∼50% (47-54%) of the cells identified on our spatial transcriptomics. While existing scRNA datasets of arteries in the past have demonstrated inflammatory cells being the most numerous populations of isolated cells^28,38,39^, our more vigorous tissue digestion protocol allowed for identification and isolation of SMC as by far the most numerous cell type, matching histological observations. SMC were classically identified using expression of mature smooth muscle contractile apparatus genes such as SMC heavy chain (*MYH11*) or calponin 1 (*CNN1*) (Figures 2A and 2B). Visualizing the population using several canonical SMC marker genes revealed a continuous gradient of SMC maturity in healthy vessels as marked by a gradient of expression of each of these “SMC-specific” markers. SMCs (as evidenced by RNAscope and spatial transcriptome), exist in a distinct gradient of hierarchical cell-states, *MYOCD*^+^>*CNN1*^+^>*MYH11*^+^>*ACTA2*^+^>*TAGLN*^+^ population. (Figure 2A). It is important to note that a subset of commonly applied smooth muscle markers (smooth muscle actin/*ACTA2*, and transgelin/*TAGLN*/SM22α) are expressed quite broadly, including very robust expression of *ACTA2* and *TAGLN* in fibroblast and a subset of endothelial cells. This has significant implications regarding lineage tracing models using *Tagln* or *Acta2* loci as the driver for Cre-recombinase lineage labeling, and perhaps explains the distinct results obtained using different Cre-recombinase in the existing literature^40^.

**Figure 2.**
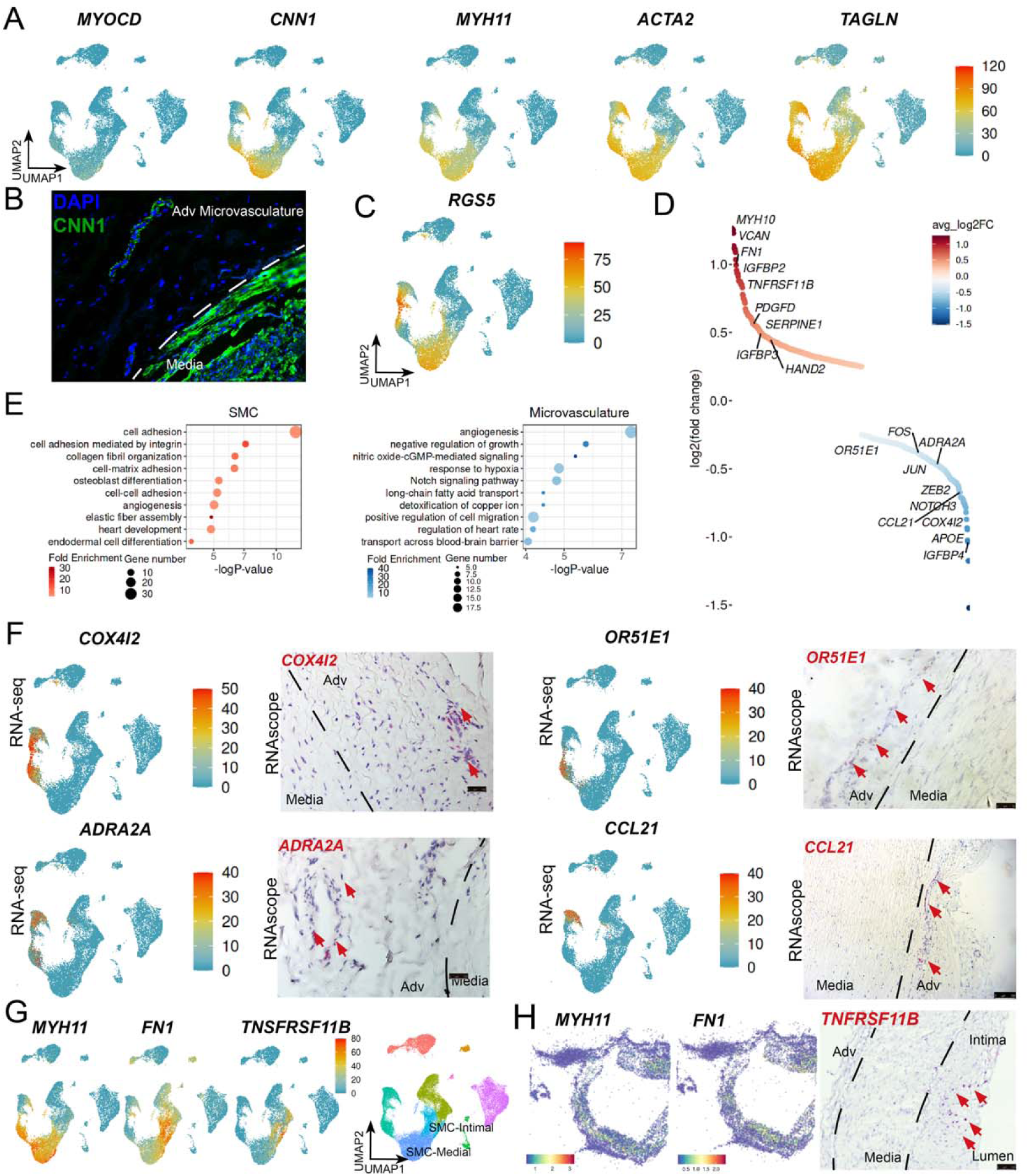
Smooth muscle cells of great arteries. (A) UMAP visualization of single-cell transcriptomes showing the expression levels of key mature contractile smooth muscle markers, including *MYOCD, CNN1, MYH11, ACTA2*, and *TAGLN*. These genes highlight the identity and maturation state of SMC populations. (B) Immunofluorescence staining of the ascending aorta tissue, visualizing the localization of CNN1 (green) as a marker of contractile SMCs, with nuclear counterstaining using DAPI (blue), confirming the spatial identity of SMCs in situ. (C) UMAP plot illustrating the expression of *RGS5*, a known marker of pericytes and a subset of SMCs, further delineating subpopulations within the smooth muscle lineage. (D) Differentially expressed genes (DEGs) ranking between the two major smooth muscle cell clusters, based on average log₂ fold change, indicating transcriptional divergence and potential functional specialization. (E) Dot plot summarizing Gene Ontology (GO) enrichment analysis of DEGs between the two SMC clusters (left: smooth muscle; right: microvasculature), with dot size representing gene set size and color indicating normalized enrichment (z-score), highlighting distinct biological pathways. (F) UMAPs showing marker gene expression specific to adventitial microvasculature cells (left panels). RNAscope in situ hybridization on artery tissue sections validated the expression of these markers at the spatial level (right panels), confirming the molecular identity of this rare vascular niche. (G) UMAP visualization of the expression of *MYH11*, *FN1*, and *TNFRSF11B*, genes associated with medial and intimal SMCs, suggesting spatial or functional heterogeneity among SMC subtypes. (H) Spatial transcriptomics maps showing the localization of cells expressing *MYH11* and *FN1* within the right coronary artery (RCA) (left and middle panels), coupled with RNAscope validation of *TNFRSF11B* expression (right panel), providing high-resolution spatial validation of gene expression patterns. Abbreviation: Adventitial (Adv) See also Figure S2 and Table S2.

The high depth of sequencing allowed for the identification of two large transcriptomically distinct populations (labelled “SMC” and “Microvascular” in Figure 1C). Both populations express canonical SMC markers *MYH11/CNN1* (Figure 2A). However, they both consist of subgroup of a more “contractile” population and a more “phenotypically modulated” population whereby the contractile markers are lost. The presence of MYH11-negative cells clustering with smooth muscle populations, rather than fibroblast or other cell types, may indicate phenotypic modulation of SMCs, a process well documented in murine lineage tracing models. Alternatively, it could be a mathematical artifact from gene-expression-based clustering. To distinguish the two possibilities, we leveraged murine scRNAseq data that contained Myh11-Cre lineage tracing^28^ to identify SMC and non-SMC progenies. Mapping the transcriptomes of these SMC progenies onto the human scRNAseq data revealed that both the “SMC” and “Microvascular” clusters correspond to murine Myh11-lineage traced cells, including those lacking contractile SMC gene expression (Figure S2A). This suggests that cellular lineage relationships may be captured through unbiased clustering methods. We subsequently performed more detailed analysis of the two SMC populations.

Staining for specific contractile SMC marker *CNN1* revealed positive cells in both the media and small vessels in adventitia (Figure 2B). We therefore hypothesize that these two populations of SMC reflect medial (larger cluster) and microvascular SMCs (smaller cluster). To better understand this observed heterogeneity of these two distinct populations of cells expressing canonical “smooth muscle markers,” we performed differentially expressed gene analysis across these two populations, which revealed 587 significantly differentially expressed genes (Figures 2D and S2B; Table S2). Microvascular mural cells / pericytes were classically identified using expression of markers *RGS5* and *NG2/CSPG4*^34,41^. Interestingly, these “canonical pericyte markers” are not confined to one transcriptomically distinct population, but rather mark non-overlapping cell populations (Figures 2C and S1A), suggesting that cells previously categorized as pericytes in different studies using these markers may represent a heterogeneous population of cells, including subsets of medial SMC. Markers more specific to the smaller population of SMC (microvascular population) include *COX4I2, IGFBP4* and *OR51E1.* Leveraging RNA in situ hybridization and the spatial transcriptome with Slideseq for markers unique for the smaller SMC population confirmed that the smaller population of cells expressing contractile markers resides exclusively in the adventitia (Figures 2F and S2E), and therefore represent mural cells of the microvasculature. While these cells do express some of the classical “pericyte” markers, due to the potential imprecise nature of these markers as described above, we will refer to this population of cells as “adventitial microvasculature” cells. While both of these clusters are MYH11+ cells in the vascular wall, these two populations of SMCs have very distinct biologies. Pathway analysis demonstrated that these two populations have distinct biological functions (Figures 2E and S2C). For example, these two populations of SMCs from the same segment have distinct patterns of adrenergic receptor expression with alpha adrenergic receptor 2A (*ADRA2A*) (Figure 2F) found only in the microvascular population, whereas *ADRA2C* is found more in the medial SMC (Figure S2D).

Given the diversity of transcriptome that was observed in the microvascular population, we performed further sub-clustering of this population into two distinct clusters. Transcriptomically, these two clusters have unique markers including *CCL21*, *OR51E1* (Figures S2F and S2G; Table S2). Spatially both of these clusters reside in the adventitia by RNAscope and Slideseq (Figures 2F and S2H). Pathway analysis revealed these two population of microvascular cells likely also have distinct function, with one population more enriched for “vascular development”, while the other more enriched for “signal transduction” and “cellular migration” processes. These data suggest at least two or more populations of microvascular cells exist in the adventitia with distinct function.

We then turn our attention to the largest SMC cluster, which is the most abundant cell type in the arteries and maps to the vascular media and intima. This population of SMC can be further divided into two subclusters whereby one subcluster express high level of mature SMC markers, and another subcluster where the expression of contractile genes such as *MYH11* or *CNN1* are absent (Table S2). To identify where these cells were located, we performed spatial transcriptomic profiling using Slideseq in conjunction with RNAscope for specific markers for this population. Consistent with previous work^28,42^, one of the most specifically expressed genes in these SMC subclusters included *TNFRSF11B* and *FN1*. The less “contractile SMC” cells localized to the intima (Figures 2G, 2H and S2I). Again leveraging existing murine scRNAseq data with lineage tracing, we found that the less contractile subcluster of SMC transcriptomically map to “phenotyptically modulated SMC” in Myh11 lineage tracing models. Previously identified markers for murine modulated SMC^28^ are found in the same intimal cluster (Figure S2J), including rare cells that express “chondrogenic SMC” markers such as *IBSP1* and *HAPLN1* (Figure S2K). Applying lineage inference tool using pseudotime, a similar de-differentiation timeline can be constructed using human medial/intimal SMC (Figure S2L). These findings suggest that the majority of the cells that makes up the human healthy arterial intima shares significant transcriptomic similarity to phenotypically modulated SMC observed in murine models (Figure S2A). The human data, in conjunction with *Myh11*-cre SMC lineage tracing in murine models^28,42^, lend further evidence to the notion that arterial intima, which is present in human but not mice, are likely smooth muscle derived, arising from media SMC in a process analogous to neointimal formation in murine atherosclerosis and intimal hyperplasia models.

### Adventitial Fibroblast in Human Arteries

The second most abundant population of cells isolated in our dataset are adventitial fibroblasts, which form a continuum in the transcriptomic space with smooth muscle cells. This population of cells did not express contractile SMC markers (Figures 1C and 2A), but did express high levels of extracellular matrix components and modifiers (Figure 3A). Marker gene RNAscope studies for this population, along with spatial transcriptomics, localized this population exclusively to the adventitia (Figures 3B and 1D). Based on marker gene expression, we termed this population vascular adventitial fibroblast.

**Figure 3.**
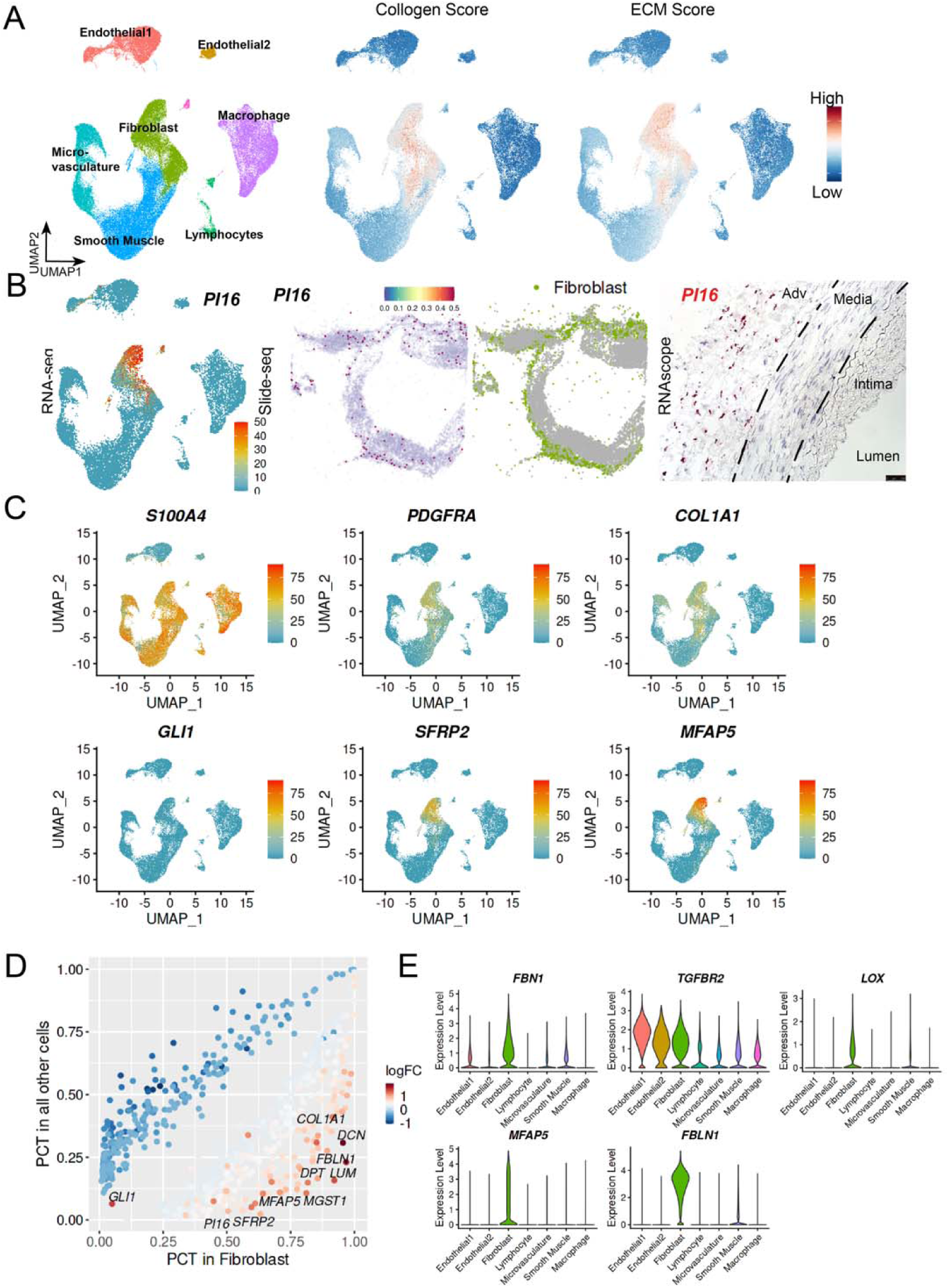
Adventitial fibroblast in human arteries. (A) UMAP plots showing composite scores derived from the expression of collagen-related genes (middle) and broader extracellular matrix (ECM) gene signatures (right), highlighting their enrichment in specific cell populations. Cell type annotations are shown on the left for reference, indicating that fibroblasts are the predominant ECM-producing cells. (B) *PI16* RNA expression visualized on the RNA-seq UMAP (left) and Slide-seq spatial transcriptomics map (middle), revealing specific enrichment of PI16-positive fibroblasts in the adventitial region of the right coronary artery (RCA). This localization was further validated by RNAscope in situ hybridization using gene-specific probes (right), confirming the spatial identity of PI16-expressing fibroblasts. (C) UMAP visualization showing RNA expression levels of canonical fibroblast marker genes, including S100A4, PDGFRA, COL1A1, GLI1, compared to new markers SFRP2, and MFAP5. (D) Differential gene expression analysis between fibroblasts and all other arterial cell types. Each gene is plotted by its average log_2_ fold change and the percentage of cells expressing it, highlighting fibroblast-enriched transcripts involved in extracellular matrix organization and tissue remodeling. (E) Violin plots showing the selective expression of key adventitial fibroblast genes, including *FBN1, TGFBR2, LOX, FBLN1*, and *MFAP5*, which are associated with matrix assembly, TGF-β signaling, and vascular structural integrity. These markers reinforce the functional identity of fibroblasts as essential ECM-regulatory cells within the arterial wall. Abbreviation: Adventitial (Adv) See also Figure S3 and Table S3.

Commonly used canonical fibroblast marker genes^32,43^, including fibroblast specific protein (FSP1/*S100A4*), *PDGFRA*, *COL1A1*, *GLI1,* were expressed relatively diffusely in several different populations of non-fibroblast cells (Figure 3C). This likely explains part of the heterogeneity of results that comes from different fibroblast lineage tracing / manipulation Cre models in murine studies^44–46^. Using the scRNAseq data, we additionally identified *PI16*, *SFRP2*, and *MFAP5*as the most specific marker genes for this adventitial fibroblast population (Figures 3C and 3D; Table S3). Of note, genes highly expressed specifically in the adventitial fibroblast population were enriched for genes known to cause genetic aortopathies^47–49^, including *FBN1*, *TGFBR2*, *LOX*, *FBLN1*, and *MFAP5* (Figure 3E). This suggests a potentially prominent role for this population of cells in the etiology of aortopathies and other vasculopathies.

Surprisingly, while it has been described that the adventitial fibroblast is a heterogeneous population, they appeared to be relatively homogenous in transcriptomic space within the same vascular segments. There were additional endothelial cells and other microvascular cells that are in the adventitia, which makes it more heterogenous than other layers, but the fibroblast themselves within each segment appeared to be mostly one large continuous population across a spectrum. However, there did appear to be significant heterogeneity across the adventitial fibroblasts among the different vascular beds (Figure S3A). Focusing on the transcriptomic heterogeneity across the adventitial fibroblast population, three distinct clusters highlighted by different biological processes can be found (Figures S3B and S3C). Pathway analysis of differentially expressed genes across these clusters revealed particular sensitivity of each of these clusters to different signaling pathway (Figure S3D). One cluster has particular enrichment in Fibroblast Growth Factor and pro-migration signal (we termed “FGF-avid”), another has higher TGFB pathway and ossification signals (we termed “TGFB-avid”), and the third has enrichment for BMP signals and microfilaments such as MFAP5 and FBN1 (we termed “BMP-avid”) cluster (Figures S3C, S3D and S3E). While fibroblast carrying each of these subsets exist in each vascular segments, the “TGFB-avid” fibroblast are particularly enriched in the root, ascending, and arch, and PA, corresponding to those with neural crest origin, whereas “BMP-avid” cluster is particularly enriched in the descending, infra- renal, and iliac arteries, corresponding to those with paraxial/splenic mesodermal origin (Figure S3A).

### Endothelial Cells

Endothelial cells heterogeneity across different types of vessels has been well described, including at the single cell level in the organ specific endothelial atlases^35,50,51^. Even at a specific arterial site, there is evidence of different populations of endothelial cells as determined by single cell RNA sequencing of murine aorta^35^. Well established markers for endothelial cells (*CDH5*, *VWF, CD31*) fall within two large distinct clusters (Figures 4A and S4A). A single layer of endothelial cells is found in the intimal surface of the arteries, but frequently many more are found in the adventitia associated with the adventitial micro-vasculature. In our data, the microvascular endothelial cells appear to be the most abundant population of endothelial cells (Figure 4B). The biggest determinant of the transcriptome of the endothelial cells appears to be which region of the vascular wall (intima/adventitia) these cells occupy (Figure S4B). The larger of the two clusters constitutes the majority of the endothelial cells (“Endo1”), and can be further subdivided into additional subclusters (Figure 4C). One subcluster expresses markers previously found to be associated with venous and capillary endothelium, (for example, *ACKR1*), whereas another subcluster expresses *HEY1*, a Notch signaling responsive gene previously associated with arterial endothelium (Figure 4D). Consistent with existing literature, the cluster that expresses *ACKR1* is found exclusively in the microvasculature in the adventitia, whereas the cluster that expresses *HEY1*, a key gene associated with arterial endothelial specification^52^, is found both in the adventitia as well as on the luminal surfaces. These data suggest that these EC reside in arterioles in the adventitia as well as on the vascular surfaces (Figures 4E and S4C). Arterial endothelial cells have been implicated in endothelial to mesenchymal transition (EndoMT), whereby these cells start to express mesenchymal genes^53,54^. We did find rare endothelial cell populations that express *TAGLN* and *ACTA2,* but total incidence is (0.57% total cells), which is below the doublet rate. While these cells are not flagged by computational doublet removal tools^55^, we cannot entirely eliminate that these may simply be doublets (Figure S4D). Lastly, using existing markers for lymphatic endothelial cells (*PROX1*, *LYVE1*)^56,57^, we can observe a small population in the adventitial EC as well, though they consist of a very small minority (Figure 4F).

**Figure 4.**
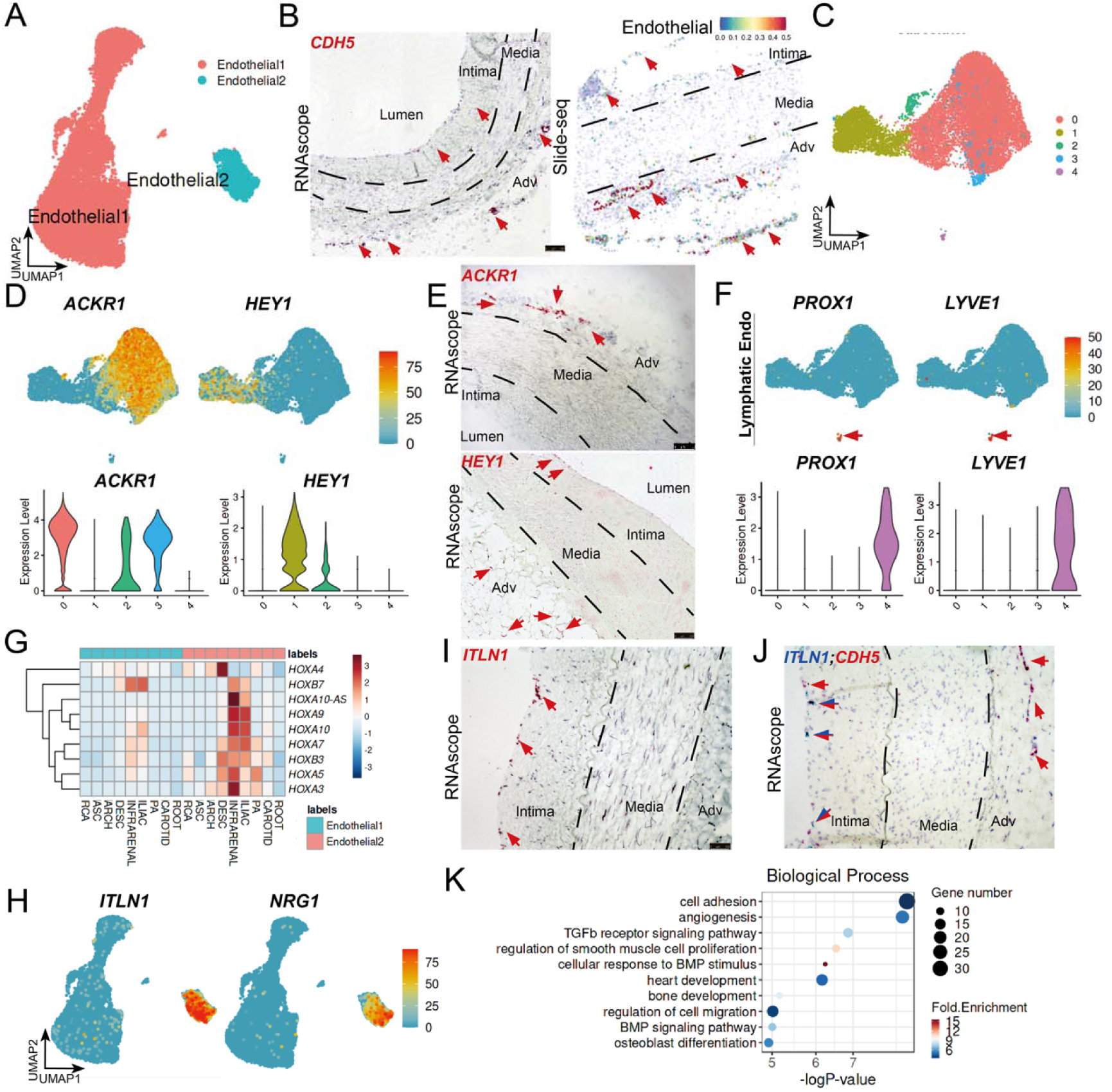
Distinct endothelial cell populations across vessels. (A) UMAP visualization of single-cell transcriptomes revealing two major, transcriptionally distinct clusters of endothelial cells, indicating the presence of heterogeneous endothelial subpopulations within the human arterial system. (B) Validation of endothelial cell identity using RNAscope in situ hybridization for the canonical marker *CDH5* in the ascending aorta (left), alongside spatial transcriptomics (Slide-seq) mapping of endothelial cells in the aortic wall (right), confirming their vascular localization. (C) Sub-clustering analysis of the Endothelial 1 population, revealing finer substructures suggestive of functional or anatomical specialization within this cluster. (D) UMAP plots (top) and violin plots (bottom) depicting the expression of representative cluster- specific genes within Endothelial 1 cells: *ACKR1* (left) and *HEY1* (right), indicating divergent functional programs among subpopulations. (E) Spatial validation of *ACKR1* (top) and *HEY1* (bottom) expression in the ascending aorta using RNAscope probes, confirming the localization of transcriptionally distinct endothelial subtypes in situ. (F) UMAP (top) and violin plots (bottom) showing the expression of non-endothelial lineage genes within Endothelial 1 cells: *PROX1* and *LYVE1*. (G) Heatmap showing the expression z-scores of *HOX* genes across different vessel segments, stratified by Endothelial 1 and Endothelial 2 clusters. The distinct expression patterns reflect segmental identity and positional memory encoded by *HOX* gene signatures. (H) UMAP plots displaying expression of Endothelial 2-specific genes *ITLN1* and *NRG1*, highlighting their restricted expression pattern and suggesting specialized endothelial functions. (I) RNAscope staining of *ITLN1* expression in the ascending aorta, validating the RNA-seq findings and confirming spatial localization of the Endothelial 2 subtype. (J) Dual RNAscope staining of *ITLN1* (blue) and *CDH5* (red) in the ascending aorta, demonstrating co-localization of general and subtype-specific endothelial markers, with double positive cells found only on the luminal/intimal endothelium. (K) Dot plot showing the enrichment of biological processes based on differentially expressed genes between Endothelial 1 and Endothelial 2. Dot size reflects gene set size, and color indicates normalized enrichment scores (z-scores), emphasizing functional divergence between these arterial endothelial subtypes. Abbreviation: Adventitial (Adv) See also Figure S4 and Table S4.

The second cluster of endothelial cells (“Endo2”) are notably transcriptomically distinct from the remaining endothelial cells. Interestingly, even within the same segment, this population of cells appears to express different *HOX* genes compared to the other larger endothelial population, suggesting they likely come from distinct origins (Figure 4G). This endothelial cell population expresses a large number of distinct markers, many with known biological function and vascular disease relevance, including *ITLN1*, *NRG1*^58–60^ (Figure 4H). The Endo2 population is found in all the different vascular territories, and accounts for a ∼10% of the total endothelial population and does not appear to significantly vary in proportion across different vascular beds (Figure S4E). Interestingly, one of the most specific markers, *ITLN1*, is known to be upregulated after large vascular ischemic events^61–63^. RNAscope using *ITLN1* revealed that this population exists exclusively on the luminal surface, and are not found in the adventitia similar to the other endothelial (Endo1) population (Figures 4I and 4J). Compared to previously reported data, the marker gene for this population appears distinct from the different endothelial populations reported in murine aortic endothelial cells^35,51^. However, the same distinct population of endothelial cells appears to be present in other human large artery single cell datasets^38,64^. Differentially expressed (DE) gene comparison between the two endothelial cell populations (Figure S4F; Table S4) revealed that they likely have distinct functions.

Compared to the numerically more abundant Endo1 population, the Endo2 population enriches for genes associated with TGFb/BMP signaling, angiogenesis and SMC crosstalk, as well as a strong disease association with atherosclerosis (Figures 4K and S4G). These findings suggest this Endo2 population may be uniquely involved in human vascular diseases.

### Macrophages And Lymphocytes

While classically not considered part of the vessel wall, emerging evidence suggests that inflammatory cells play an important part in vascular homeostasis^11,33,64–66^. Our data further supports this notion, as even though these donor samples were obtained from relatively healthy vessels without clinically overt diseases, there is a significant number of macrophages and lymphocytes present in all the vascular beds. There is a trend towards a slight increase in the total fraction of immune cells in the atherosclerosis prone arterial beds (coronary, infra-renal aorta) (Figure S5A) but does not reach statistical significance. There was also no significant inter-person difference in proportion of immune cells. (Figure S1E) The majority of the macrophages and lymphocytes are found in the adventitia. Clusters of B and T cells can also occasionally be found in organized centers of what appears to be tertiary lymphoid organs that abut the adventitia (Figure 1D). Putative computational analysis of cell-to-cell crosstalk using CellChat^67^ demonstrates significant cross talk between the adventitial fibroblast, smooth muscle, and the immune cell populations (Figure S5B). These findings suggest a potential role for resident vascular mural cells in coordinating the vascular immune response.

### Determinants of Arterial Identity

To understand differences between arterial segments, we first tried to determine whether there was a consistent relative composition of different cell types within different arteries. Using the low clustering resolution which identified the canonical populations of vascular cells, we discovered that with identical digestion protocols, the relative proportion of cell populations were mostly similar across different arterial beds (Figure S6A). There is a slight increase in inflammatory cells in the vascular beds that are more prone to atherosclerosis, such as the coronary artery, and infra-renal segment of the aorta (Figure 5A). However, there were no large differences in cellular composition across arterial segments, similar to histological studies.

**Figure 5.**
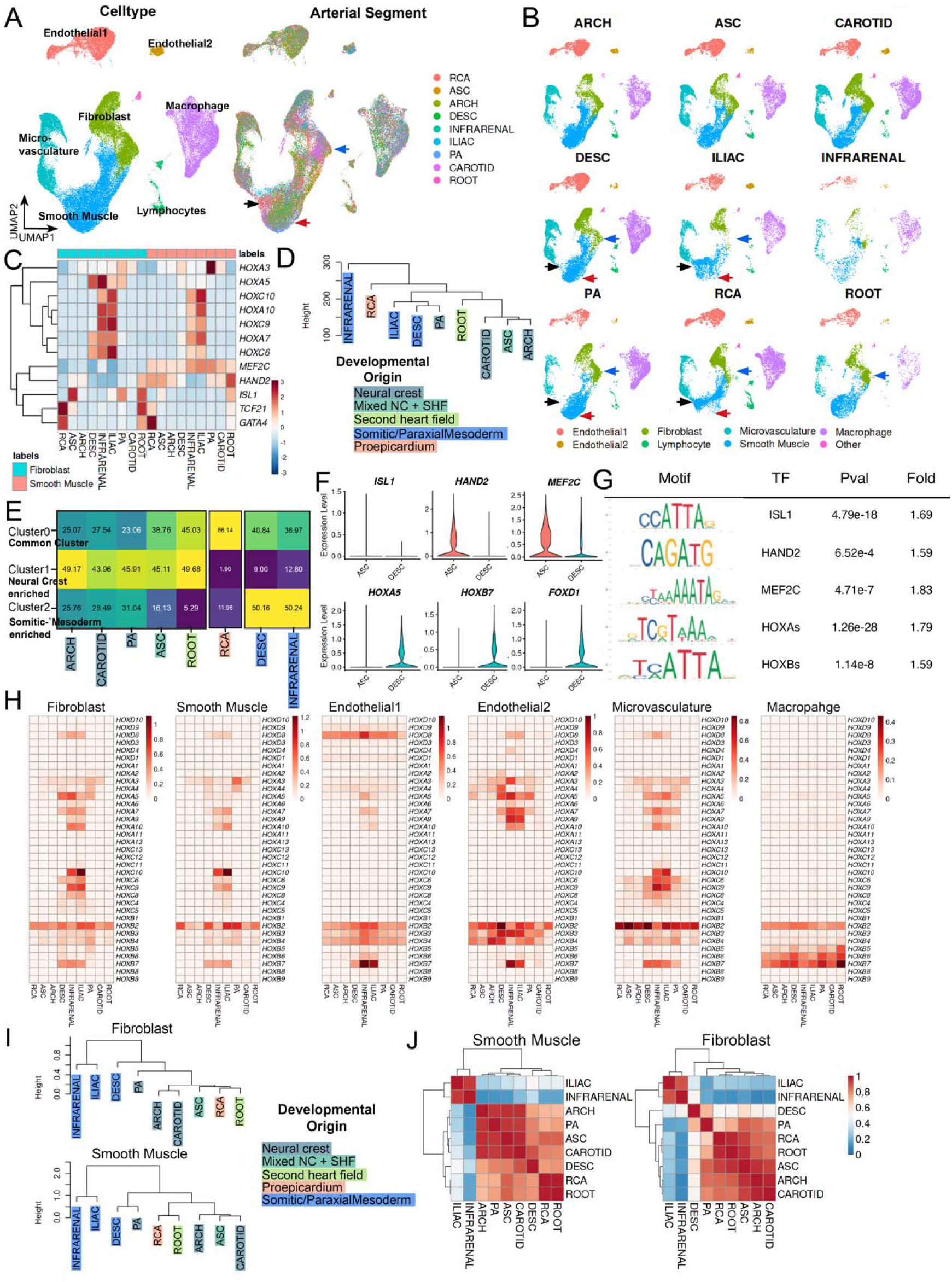
Prominent roles of embryonic origin in adult vascular identity. (A) UMAP visualization showing the transcriptional heterogeneity of major vascular cell populations—smooth muscle cells (SMCs), fibroblasts, endothelial cells, microvascular cells, and macrophages—across distinct vascular beds, indicating segment-specific cell states. (B) Split-view UMAP plots displaying the segmental origins of each cell, revealing that within each major cell type, transcriptomic profiles vary substantially across different arterial regions, underscoring spatially encoded identities. (C) Heatmap showing the z-score–normalized expression of *HOX* genes across multiple vascular segments, in both SMCs and fibroblasts. The segment-specific expression patterns indicate the preservation of developmental *HOX* code into adult vasculature. (D) Hierarchical dendrogram clustering of fibroblast transcriptomes across vascular segments, using *Euclidean* distances computed from global gene expression profiles. Segments are grouped according to transcriptional similarity, which aligns closely with known embryonic origins (e.g., neural crest vs. mesoderm). (E) Heatmap quantifying the relative representation (percentage) of each arterial segment within transcriptionally defined UMAP clusters, highlighting regional bias and the compartmentalization of cell states. (F) Violin plots depicting the expression of key transcription factors and developmental regulators—*ISL1, HAND2, MEF2C*, and representative *HOX* genes—in fibroblasts from ascending versus descending vascular regions. These patterns support region-specific regulatory programs. (G) Transcription factor binding motif enrichment analysis showing overrepresentation of ISL1, HAND2, MEF2C, and HOX motifs in the promoter regions of differentially expressed genes between ascending and descending fibroblasts. Macrophages were used as a negative control, demonstrating fibroblast-specific enrichment. (H) Heatmap showing log_10_-transformed normalized expression values of *HOX* genes across multiple vascular segments in six major cell types: fibroblasts, SMCs, Endothelial 1, Endothelial 2, microvascular cells, and macrophages. The distribution reveals cell type– and segment– specific *HOX* gene signatures. (I) Dendrograms displaying hierarchical clustering of vascular segments in fibroblasts (top) and SMCs (bottom), based on the expression of *HOX* genes. Clusters reflect developmental lineage relationships and segmental positional memory. (J) Pearson correlation heatmap showing inter-segmental transcriptional similarity based on *HOX* gene expression in SMCs (left) and fibroblasts (right). Segments are organized by Euclidean distance clustering, revealing conserved spatial transcriptional identities within and across lineages. Abbreviations: ascending thoracic aorta (ASC), aortic arch (ARCH), descending thoracic aorta (DESC), infra-renal abdominal aorta (INFRARENAL), iliac artery (ILIAC), coronary artery (RCA), carotid artery (CAROTID), pulmonary artery (PA), and aortic root (ROOT), Neural crest (NC), Second heart field (SHF) See also Figure S7 and S8; Table S5.

Visualization of a large dataset using UMAP accentuates the drivers of the variance within the dataset^68,69^. Thus, visualization of the data in this manner also enables us to get a sense of relative heterogeneity of each population of cells across the different vascular beds. Simply visualizing this dataset coded by segmental origins, it became readily apparent that there is significant heterogeneity between adventitial fibroblasts and smooth muscle cells among the different vascular beds. However, there were relatively fewer differences between the microvascular, inflammatory and endothelial populations across different vessel beds, as demonstrated on the UMAP (Figure 5B). Because dimensional reduction such as UMAP can distort true distances between populations^68,70^, we repeated the analysis by looking at the number of differentially expressed genes between the ascending and descending arteries, and between the coronary and descending pairs (Figures S6B and S6C), as well as the mean Euclidean distance^71^ between the average global gene expression of each cell population across the vascular beds (Figures S6D and S6E). Consistent with the UMAP visualization, both of these analyses revealed fewer differences between endothelial cells and macrophages across different vascular beds compared to fibroblast and smooth muscle cells. Interestingly, the same analysis comparing Endo1 and Endo2 population showed much smaller difference between Endo1 population but significantly more difference in the Endo2 population across vascular beds (Figure S6E). These data suggest that the main determinant of arterial identities, at least transcriptomically, appears to be held mostly by fibroblast, smooth muscle cells, and possibly the smaller subset of intimal endothelial cells, Endo2 population, and much less so in the remaining endothelial cells, microvasculature, and inflammatory cells.

### Prominent Roles of Embryonic Origin in Adult Vascular Identity

We next investigated the main differences between SMC and fibroblasts from different vascular beds. Examination of the differentially expressed genes across SMC and Fibroblasts of different vascular beds revealed a large number of developmentally important transcription factors^28,72–77^, including *HOX* family genes, *ISL1*, *MEF2C*, *TCF21, HAND2, GATA4* etc. (Figure 5C). We hypothesize, therefore, that embryonic origins are the dominant factor in vascular identity. We regenerated a UMAP visualization focusing on the top transcriptomic variances that exist in each individual population. This UMAP readily revealed distinct transcriptomic sub-clusters of Fibroblast and SMC (Figures S3B and S7A). Consistent with our hypothesis, the proximity of different populations of fibroblast by either Euclidean distance or visualization in UMAP space appears to correspond to each segments presumed embryonic origin reported in animal models (Figures 5D, 5E and S7B). Cells from arteries of different caliber and flow with similar embryonic origins (i.e.aortic arch, and ascending aorta/carotid arteries) cluster closer together than their connected counterparts (such as descending thoracic artery to aortic arch). This suggests that embryonic origins of each of these arterial segments play a dominant role in adult gene expression. Curiously, this developmental pattern is less preserved in identical analysis of macrophage, microvasculature, and Endo1, suggesting that in these cells the resting gene expression may be less influenced by their embryonic origin (Figures S7C, S7D and S7E).

Many of the differentially regulated genes found on the list has been known to be regulated by these transcription factors in different processes. To unbiasedly determine if these lowly expressed developmental master-regulators continue to play an important role in adult gene expression globally, we investigated whether the binding sites for these transcription factors are enriched in differentially expressed genes between segments compared to other variable genes in the dataset, focusing on the ascending and descending thoracic aorta as the main test case. Transcription factors *ISL1*, *HAND2*, *MEF2C* are expressed in the second heart field and neural crest cells which give rise to the ascending aorta, whereas *HOX* genes are expressed in the somitic mesoderm which gives rise to the descending thoracic aorta (Figures 5F and S7F). Consistent with these transcription factors playing an important role, we saw a very strong enrichment of ISL1, HAND2, MEF2C, and HOX binding sites in the regulatory regions of differentially expressed genes in fibroblast and SMC across these vessel beds of distinct embryonic origin (Figures 5G and S7G; Table S5). These data suggest the embryonic origin and the persistent expression of these critical transcription factors into adulthood plays a critical role in arterial identity and their global gene expression in adult.

### Adult Vascular Hox Gene Expression Patterns

While *HOX* genes’ roles are best elucidated in development, as one of the most potent cell fate and patterning signals^72,78^, their role in adult tissue identity and disease is less well understood. It has been shown that different arterial segments express different *HOX* genes by bulk RNAseq^25^. Given the impressive enrichment of *HOX* gene binding motifs in regulatory regions of most differentially expressed genes, we leveraged our dataset to better understand *HOX* gene expression across the different cell types within each segment of the aorta.

It appears that adult *HOX* gene expression patterns from the same vessel segment are conserved in adulthood and thus mirror their embryonic origin (Figures 5H and S8A). Descending thoracic aorta, abdominal aorta, and iliac arteries each expresses distinct subsets of *HOX* genes that are conserved from one person to the next (Figure S8B). However, we observed that within the different cell types of a segment, each cell type expresses a different subset of *HOX* genes. For example, in descending thoracic aorta, *HOXD8* is most highly expressed only by Endo1, *HOX A4/A5* predominantly by Endo2, and *HOXA3* in SMC, and *HOX A5/B7* in fibroblast. While smooth muscle and microvasculature are transcriptionally and phenotypically similar, and AdvFib and microvasculature are geographically close, they express very distinct *HOX* genes patterns (Figures 5H and S8C). On the other hand, adventitial fibroblast *HOX* gene expression is quite similar to those of the medial SMC. This potentially reflects these two populations of cells (medial SMC and AdvFib) have more closely shared embryonic origins compared to those of microvasculature in the adventitia. As further evidence that the *HOX* genes likely play an important role in adult arterial identity, we clustered the cells and segments based on *HOX* genes alone. Hierarchical clustering of SMC and fibroblast based on *HOX* genes alone mirrors hierarchical clustering of whole transcriptomes, and readily reflects each segment’s developmental origin (Figures 5I and 5J). Given that most immune cells are likely formed uniformly in the marrow and migrate into each segment, unsurprisingly there is less distinction between them. While the endothelial cells are some of the earliest to form during embryonic development of the great vessels, with each segments’ endothelial cells having distinct origins, they are significantly less heterogeneous compared to neighboring segments of different embryonic origin, with the potential exception of the Endo2 population (Figure S8D).

Those populations that differ from one vascular bed to another tend to have a very distinct pattern of *HOX* gene expression, whereas cell types with similar *HOX* gene expressions tend to have globally more similar transcriptomes. The persistence of this very distinct pattern of *HOX* gene expression across individuals suggests that conserved transcriptomic regulation of these genes in adulthood, along with enrichment in HOX binding motifs in differentially expressed genes noted above suggests critical roles for *HOX* genes in cell type specific adult segmental specific gene expression.

### Enrichment of Vascular Disease Rare and Common Variant Genetic Signals in Segmental Transcriptomic Differences

Our original hypothesis was that the transcriptomic differences between these vascular beds are important for differences in disease propensities, and that the single cell atlas will help identify novel pathologic populations and genetic programs. To determine if these identified transcriptomic differences play a role in adult diseases, we then asked whether the differentially regulated genes were enriched in human signals of genetic diseases. We again focused our attention on the difference between ascending and descending thoracic aortic segments, which have a known strong propensity for aortic aneurysms^79,80^. While these disease segments are subject to changes in flow, there was relatively few differences between the global endothelial cell expression of cells from these two vessel beds (Figure S9A; Table S6). However, there were significantly more transcriptomic differences between fibroblast and SMC from these different connected arterial segments. Most notably, there were a large number of differentially regulated genes in the fibroblast population that have known roles in inherited aortopathies^81^. A number of mendelian aortopathies involve haploinsufficiency of critical ECM components, including fibrilin 1 (*FBN1*), lysyl oxidase (*LOX*), microfibril associated protein 4 (*MFAP4*), microfibril associated protein 5 (*MFAP5*), and collagen 3A1 (*COL3A1*). Loss of function alleles in these genes are associated with thoracic aortic aneurysms, with a strong preference for affecting the root and ascending aorta. Interestingly, the expression of each of these genes is significantly lower in the ascending than the descending aorta (Figures 6A and S9A). Closer examination revealed that differences in expression were seen nearly exclusively in the adventitial fibroblasts, and not in SMC (Figure 6B). Conversely, *PRKG1* is a gene whose gain of function is known to increase the risk of aneurysm^82–84^, also has higher expression in the aortic root and ascending compared to descending (Figure 6C).

**Figure 6.**
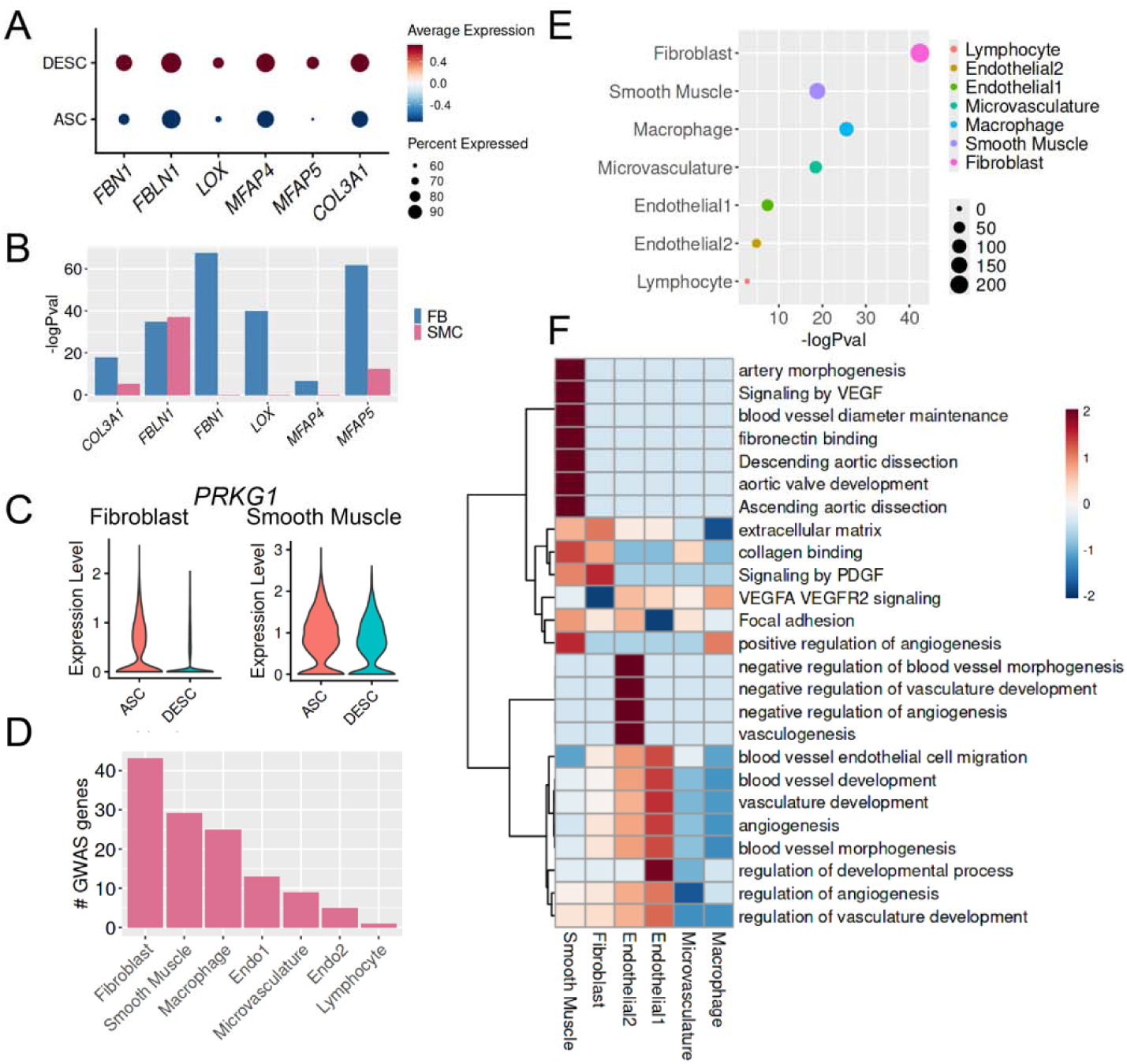
Enrichment of vascular disease variants in segmental transcriptomic differences. (A) Dot plot depicting the expression patterns of Mendelian aortopathy-related genes—*FBN1, FBNL1, LOX, MFAP4, MFAP5*, and *COL3A1*—in fibroblasts derived from ascending (ASC) and descending (DESC) aortic regions. Dot size reflects the percentage of expressing cells, and color represents scaled average expression levels, revealing regionally enriched gene programs. (B) Bar plot summarizing the statistical significance of expression differences for the same set of Mendelian aortopathy genes between fibroblasts (FB) and smooth muscle cells (SMC) from ascending and descending aortic sites, highlighting key regulators associated with thoracic aortic disease. (C) Violin plots showing the expression levels of selected aneurysm-risk genes in fibroblasts (left) and smooth muscle cells (right), stratified by ascending versus descending vascular origins. The distinct expression patterns suggest site-specific vulnerability to aneurysmal remodeling. (D) Bar graph showing the number of differentially expressed genes in each cell type that overlap with genome-wide significant loci identified in the Million Veteran Program (MVP) genome-wide association studies (GWAS). Comparisons were made between coronary artery or carotid segments versus the descending aorta, reflecting spatially distinct genetic risk profiles. (E) Summary plot ranking the genetic overlap between MVP GWAS hits and differentially expressed genes across coronary, carotid, and descending arterial segments within each major cell type. Bars represent the number of overlapping genes, suggesting potential cell type– and region–specific contributions to vascular disease susceptibility. (F) Heatmap displaying z-score–normalized fold enrichment of Gene Ontology biological processes in differentially expressed genes across coronary artery, carotid, and descending segments, stratified by cell type. Enriched pathways highlight functional divergences driven by both anatomical location and cellular identity. See also Figure S9 and Table S6, S7, S8.

To determine if other arterial segment specific vascular traits showed segregation of genetic signals, we subsequently performed the same analysis on common genetic variants associated with human aortic diameter. Most GWAS lead variants are located in non-coding regions of the genome, likely exerting its effect on gene dosages. Thus, GWAS “hits” enrich for genes whose dose are likely to contribute to pathology, with the added benefit that GWAS “hits” do not hold bias towards known pathways. We leveraged recently published GWAS on aortic diameter^85^, and asked how often GWAS genes for aortic diameter are differentially expressed in different populations of cells in aneurysm prone (i.e. ascending) vs resistant (i.e. descending) segments. For those within the non-coding region of the genome, we used the putative target gene suggested in the original manuscript, which mostly reflected the closest gene. From the list of GWAS genes, we then determined how often these dose sensitive genes were differentially expressed at the individual cell level in ascending and descending aorta. After Bonferroni correction for multiple hypothesis testing, our analysis revealed 31 diameter sensitive genes are differentially expressed in ascending vs descending fibroblasts, and 18 diameter sensitive genes are differentially expressed in smooth muscle cells, and 5, 3, and 4 genes are differentially expressed in microvasculature, macrophage, and endothelial cells, respectively (Table S7). These analyses suggest a large proportion of segment-specific modifiers of aortic propensity for aneurysm act through fibroblast and SMC, with surprisingly little transcriptomic differences in the endothelial cells (Figures S9B and S9C). These findings confirm the known role of SMC in aneurysms, while suggesting a prominent unrecognized role of AdvFib in the same disease process.

We subsequently performed the same analysis in atherosclerotic disease and observed a similar pattern. Coronary artery and carotid arteries are known to be more prone to atherosclerosis whereas descending thoracic aorta is less so^86^. Thus, these vessel beds are used for the atherosclerosis disease propensity comparisons to identify causal cells and causal genes for the reasons for the differences in disease propensity. A significant subset of common variant associated genes for coronary artery disease (Figures 6D and 6E) and aortic aneurysm (Figures S9D and S9E). are differentially expressed in the different population of cells in the vessel wall. Our analysis revealed 45 CAD GWAS genes differentially expressed in fibroblast, 30 in SMC, 11 in microvasculature, 15 in Endo1, 7 in Endo2, and 27 in macrophages (Table S8). Functional analysis of enriched genes in each population of cells revealed angiogenesis related biological processes are significantly enriched in SMC, and PDGF and VEGF signaling enriched in the fibroblast population (Figure 6F). These analyses suggest that segment specific differences in vascular disease propensities appears to be mostly encoded in fibroblast and SMC, and much less so in endothelial cells.

### Sex-specific differences

At a population level, there is a distinct difference in the susceptibility for vascular diseases between patients of different biological sex^87–90^. Coronary artery disease and Marfan syndrome, for example, are known to have worse vascular manifestation in men compared to women^91,92^, whereas Takayasu arteritis or spontaneous coronary artery dissections are much more common in women than men^93,94^. The underlying cause of these sex specific differences is poorly understood. While many of the causes of the sex specific differences may be related to numerous systemic and other confounding factors that are unrelated to the cell biology at the vessel wall, this dataset containing an equal number of men versus women, offers an opportunity to explore potential differences between male and female arteries. We performed differential expression analysis comparing different cellular populations in our dataset. This analysis revealed different cell-type specific transcriptomes between male and female sex.

Unsurprisingly, many sex chromosome specific genes, such as *XIST*, were differentially regulated in all cells. The analysis revealed 119, 176, 201, 79, 196, 281, 172 sex-specific differentially expressed (DE) genes between SMC, fibroblast, macrophage, Endo1, microvasculature, Endo2, and lymphocytes respectively (Table S9). Unlike cross-segmental identities that appears to lie more in fibroblast and SMC, sex-specific differences were seen across all cell types and are more profound in endothelial and immune cells. This parallels the observations of sex specific differences observed in incidence of vasculitis and vascular endothelial dysfunction. Sex differences in different populations appears to govern different aspects of vascular biology and different diseases. Pathway analysis with the GO biological process performed on each of these DE genes revealed distinct biological process enrichments (Table S10). Different cell type DE gene overlap with OMIM data revealed distinct vascular disease associations with known sex preferences (i.e. SMC association with Ehler-Danlos, and EC association with macular degeneration) (Table S11).

### Cell type- and Segment- Specific Non-coding Transcriptome

While existing analysis of scRNAseq data has mostly focused on protein coding genes, there is an increasing recognition that non-coding transcripts such as long non-coding RNAs (lncRNAs) play an important role in health and disease^98–100^. Taking advantage of the fact that a significant portion of lncRNAs is poly-adenylated and thus captured by the 10x platform, we asked whether lncRNAs also follow segmental and cell-type specific expression by aligning these sequencing data to a library of known lncRNAs. Given the library preparation of 10x does not allow for strand specific capture, one cannot readily determine if a read is from the coding or antisense strand of a transcript. To remove any potential confounding reads from protein coding genes, we leveraged a reference lncRNA transcriptome that eliminated any segments of overlap with any protein coding mRNAs. We will refer to this as the sc-lncRNA transcriptome.

This dataset contains 53,247 unique non-coding transcripts that are found in the vascular cells (compared to 18,088 protein coding genes). Despite our stringent filtering, on average, ∼1601 (median 1284) non-coding transcripts were detected in each cell, encoding an average of ∼397 (median 323) lncRNAs per cell. This represents ∼10% in number of genes or reads compared to the protein coding genome. Differential gene expression analysis of these data comparing cells across different segments found that a significant portion of the sc-lncRNA transcriptome also exhibited cell-type and segmental specific expression (Figures 7A and 7B).

**Figure 7.**
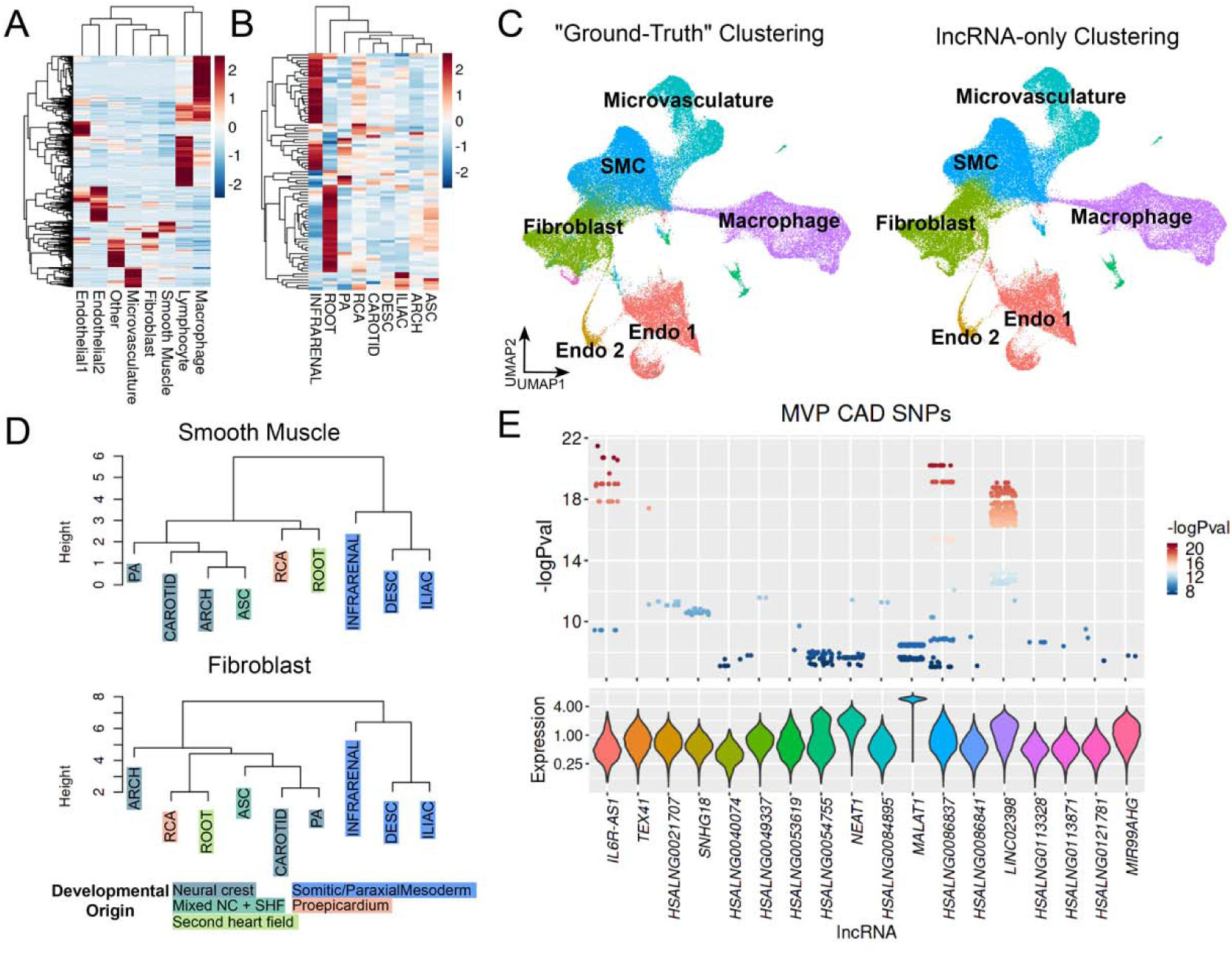
Cell type and segment specific non-coding transcriptome. (A) Heatmap showing the z-score–normalized expression profiles of long non-coding RNAs (lncRNAs) across major vascular cell types, including fibroblasts, smooth muscle cells (SMCs), endothelial cells, and immune populations. Distinct lncRNA signatures delineate cell-type– specific transcriptional programs beyond protein-coding genes. (B) Heatmap depicting the z-score–normalized expression of lncRNAs across different vascular segments, highlighting regional variation and the potential involvement of lncRNAs in positional identity. (C) UMAP projections showing single-cell clustering based solely on lncRNA transcriptomes. Left panel: clusters annotated using cell type identities derived from whole-transcriptome analysis. Right panel: clusters annotated based on unsupervised lncRNA-only clustering. The concordance and divergence between these two approaches reveal both shared and lncRNA- specific cellular features. (D) Hierarchical dendrogram clustering of arterial segments based on lncRNA expression profiles, using *Euclidean* distances in SMCs (top) and fibroblasts (bottom). Clustering patterns align with developmental origins, suggesting that lncRNAs retain positional memory reflective of embryonic lineage. (E) Top panel: Dot plot showing the overlap between MVP GWAS-identified SNPs and the loci of differentially expressed lncRNAs, indicating direct genetic association. Bottom panel: Violin plots illustrating the expression levels of these genetically associated lncRNAs, suggesting their potential role as mediators of vascular trait susceptibility. Abbreviations: ascending thoracic aorta (ASC), aortic arch (ARCH), descending thoracic aorta (DESC), infra-renal abdominal aorta (INFRARENAL), iliac artery (ILIAC), coronary artery (RCA), carotid artery (CAROTID), pulmonary artery (PA), and aortic root (ROOT), Million Veteran Program (MVP), Coronary artery diseases (CAD), Single-nucleotide polymorphism (SNP). See also Figure S10 and Table S12.

Employing these data, we investigated how much of the cell type specific identities are encoded in this limited subset of the non-coding transcriptome. To accomplish this, we performed unbiased clustering of the sc-lncRNA transcriptome using standard methods, along with UMAP based visualization. Utilizing no protein coding gene data, analyzing sc-lncRNA transcriptome alone, standard unbiased expression clustering approach could readily segregate the dataset into cell type specific clusters that recapitulate those identified using the entire transcriptome (Figure 7C). Comparison of cell type identities assigned via unsupervised low-resolution clustering to those identified by protein coding transcriptome demonstrated that ∼92% of the cells were clustered in identical ways using sc-lncRNA data alone without any protein coding transcripts. The cells exhibiting discordance were primarily situated at the interfaces between distinct cell types as visualized on the UMAP (Figure S10A). Hierarchical clustering based on sc-lncRNA data itself in SMC and fibroblast could recapitulate the developmental origin (Figure 8D). This suggests that a significant portion of the identities of each cell types, behavior, and embryological origin are embedded within the non-coding transcriptome.

The majority of common variants that are identified in humans that carry heritable risk of vascular diseases lie within the non-coding region of the genome^101–103^. One hypothesis has been that some of these non-coding regions of the genome may overlap with functional lncRNAs and alter their function. To test this hypothesis, we asked whether these vascular lncRNAs enriched for genome-wide significant GWAS SNPs for different vascular conditions. Interestingly, there appears to be a significant enrichment of GWAS SNPs for coronary artery diseases that lie within the exons of lncRNAs compared to expected (18 lncRNAs with 158 MVP SNPs) (Figure 7E; Table S12). We subsequently intersected the differentially expressed lncRNAs across segments within major cell clusters with the loci from the GWAS catalog, revealing a significant intersection of these lncRNAs with loci associated with coronary artery disease (CAD) and heart disease (Figure S10B). Repeat analysis using more updated SNPs from recent GWAS from Million Veterans Program CAD obtained similar enrichments (Figure S10C)^102^. Many of these GWAS-loci associated lncRNA have distinct expression between intimal and medial SMC populations. Intimal SMC express higher number of lncRNA transcripts than medial SMC (Figure S10D). Beyond GWAS loci, a large number of lncRNA has been associated with altered vascular disease risk^104^. Interestingly a significant portion of these previously reported vascular-disease associated lncRNA also demonstrate distinct intimal/medial SMC expression levels (Figure S10E). These findings provide potential functional context for lncRNAs implicated in vascular diseases.

## Discussion

By performing cellular transcriptomic and spatial localization of donor arteries across a variety of arterial segments, we created a resource for the vascular and genetics research community to better understand the cellular composition and distribution of gene expression programs across human arteries to better link causal genes to causal cells. We identified major cellular populations characterized not by individual markers, but by entire transcriptomic profiles and spatial localizations. We contribute to the existing literature of single cell characterization of human arteries by improving optimal digestion of highly fibrous arterial tissue to obtain high quality single cell RNAseq data superior to the cellular populations and ratios observed by histology^28,30^. These techniques also enabled us to achieve deeper characterization of vascular cell transcriptomes, including lowly expressed transcription factors, to better elucidate their biology. We hope this dataset can serve as a useful reference to aid the cardiovascular research community in future single cell characterization of vascular conditions.

Utilizing entire cellular transcriptomes instead of specific marker genes to define cellular heterogeneity, this dataset both validates and challenges existing approaches to study vascular diseases in murine and cellular models. One immediate benefit of this approach is the ability to identify specific markers empirically in an unbiased manner for populations of cells that are transcriptomically distinct. This large number of cell-type specific markers can then be utilized to enable other vascular discoveries, including generating more specific murine Cre-lox models to understand individual cell populations. Furthermore, our data can help reconcile seemingly paradoxical findings of existing cellular and lineage tracing models. For example, our analyses have revealed surprising heterogeneity and limitations of many commonly used vascular population markers (*RGS5*, *TAGLN*, *GLI1*, *ACTA2* etc), which may explain some of the contradictory findings in animal models that arise from modification using different “marker” genes^40^. This specific issue is best studied in the vascular SMC field, in which heterogeneous findings using different “SMC-specific” Cre-recombinase models have been well-documented^40^. While Acta2 and Tagln-Cre were previously frequently employed to study SMC lineage tracing and gene perturbation, meticulous studies by multiple groups have documented its activation in non-SMCs including fibroblasts^105,106^ and endothelial cells^107^. While the non-SMC labeling was previously attributed to “leaky” expression of these Cre-alleles, our scRNAseq data suggest that the issue may be underappreciated endogenous expression of these genes. Indeed, more specific SMC and vascular SMC Cres (Myh11 and Itga8^108^) have been recently widely utilized due to their specificity, which is also recapitulated in the scRNAseq data. What remains most concerning is the poor specificity of fibroblast and microvascular/pericyte markers, which is marked in our scRNAseq data as multiple populations. More dedicated markers and reagents for these populations may be needed to optimally study their biology. Similarly, by combining single cell RNAseq data with complementary spatial transcriptomic techniques, we were able to localize transcriptomically unique populations that reside in different parts of the vessel wall.

This includes identification of multiple “smooth muscle marker” positive populations, and multiple subsets of endothelial cells with distinct transcriptomes and localizations. This extensive heterogeneity that exists in healthy vessels may help explain conflicting findings that sometimes occur from in vitro studies of primary human arterial cells, as in vitro “arterial SMC” or “arterial endothelial” cells using canonical markers may be isolated from one or multiple populations of cells. Our dataset may also be used to deconvolute the population(s) of vascular endothelial cells or SMC^109,110^.

One of our main goals with this dataset was to identify underlying differences in the biology and disease propensities across different human arterial segments. Current understanding of such differences mainly attributes them to flow characteristics and hemodynamics^111–113^. Indeed, turbulent flow, laminar flow, and static flow are known to create significant alterations in transcriptomic profiles of endothelial cells that are relevant to diseases^114,115^. On the other hand, numerous pieces of evidence also exist suggesting that developmental influences may alter vascular disease propensity^116–118^, such as the frequently observed changes in atherosclerotic disease lesion quality and severity near junctions of vessels segments deriving from distinct developmental origins^119^. This nature vs nurture debate of vascular disease remains unsettled. Our analysis of healthy vessels prior to initiation of disease, yet still subjected to decades of differential flow patterns, reveals relatively few differences between the transcriptomes of endothelial cells across different vascular segments. However, adventitial fibroblasts and medial smooth muscle cells are clearly responsible for the main transcriptomic differences across different vascular beds. Differentially expressed genes in these populations of cells enrich for both common and rare variants of vascular diseases.

Furthermore, the regulatory regions of these differentially regulated genes significantly enrich for transcription factor binding sites of developmentally critical master regulators. These findings suggest that developmental origins of these vessels play a large role in different disease propensities at least equally and maybe more so than decades of differences in vascular flow and pressure. Developmental master regulators may therefore possess key pathogenic roles.

Given the clinical observation of strong associations of different arterial segments for diseases within the same person^5,80,95,120^, modulation of these developmental master regulators’ function may be sufficient to alter disease development or progression. Lessons learned from developmental biology and additional discoveries aimed to better understand these factors’ exact functions are attractive targets for future therapies.

One of the largest populations of cells identified from this dataset are adventitial fibroblasts. This population of cells carries remarkable heterogeneity across vascular segments with different disease predilections, including a large number of genetic signals for aneurysmal diseases and atherosclerosis. Yet, this population of cells has traditionally not been the focus of pathogenesis of vascular diseases. Further evaluation of this population of cells and their roles in vascular pathologies are warranted.

Analysis of the human endothelial population reveals significant heterogeneity as previously demonstrated in murine vascular single cell analysis. However, unlike murine models^35,51^, the majority of human endothelial cells in the vessel walls reside in the microvasculature of the adventitia. Consistent with distinct developmental origins and early arterial/venous/lymphatic endothelial fate commitment demonstrated in animal models^14,15,18,35,51,56,57^, the most significant determinant of the endothelial transcriptome is the type of vessel it covers. A distinct population of relatively rare endothelial cells is located at the luminal surface of the great arteries. The lack of an equivalent endothelial cell population in murine single cell vascular data suggests the function of this endothelial population may not be accurately measured in murine models. Given the enrichment of this population of cells for distinct disease-related transcriptomic programs, further dedicated studies of this population of cells may yield distinct insight into their pathophysiological function.

Murine disease models with SMC lineage tracing have identified “de-differentiated” and “phenotypically-modulated” SMC as key pathogenic cells in a variety of vascular conditions^17,28,36,37^. The importance of this population of cells is well-established through a variety of lineage tracing and manipulation models of different diseases^28,42,121,122^. However, in macroscopically healthy vessels as obtained in these studies, an apparent de-differentiated SMC subset is universally present across all vascular segments and populates the intima. This population of cells expresses a similar global transcriptomic program as those SMC-lineage traced cells that have lost their contractile genes in murine models. This potentially reflects the presence of subclinical diseases in all these donors given years of relative hyperlipidemia due to exposure to a western diet^123^. Alternatively, another explanation for this observation is that SMC phenotypic modulation itself is part of the normal physiological spectrum and not necessarily pathologic. Instead, the specific ways which seemingly similar SMC populations phenotypically modulate in different ways may govern the quality and burden of disease. This notion is further supported by the fact that there appears to be particularly large global transcriptomic differences in the “modulated” /intimal SMC population observed across the vascular beds, more so than “contractile” smooth muscle cells. This modulated SMC population also carries more genetic signals of vascular diseases than all other cell populations (Figures S9D and S9E). Existing data are unfortunately inadequate to discern these two possibilities. Additional characterization of a larger number of healthy and diseased vascular tissues at different ages may help elucidate the answer.

Lastly, while the focus of pathogenic factors for vascular disease is often on protein coding genes^85,101–103^, analysis of non-protein-coding transcriptomes of the vascular beds reveals surprisingly rich information and heterogeneity. Global analysis of lncRNAs alone can readily identify the main vascular wall cell types and differentially expressed lncRNA across vascular beds disproportionally enrich for non-coding variants that influence vascular disease risk. While the analysis is preliminary, the enrichment of genetic signals suggests these differentially enriched lncRNAs may contain critical regulators of disease risk. This finding add on to growing literature of pathogenic LncRNA in different human cardiovascular diseases^104^, and points to new potential disease modifying lncRNAs that may be targeted for cardiovascular diseases.

## ACKNOWLEDGMENTS

This work is made possible by the generosity of our tissue donors who donated their organs to save others’ lives and expand our scientific understanding of the human vasculature. This work was supported by Human Cell Atlas grant (ZF2019-002437) from the Chan Zuckerberg Foundation (T.Q., P.C); National Institutes of Health grants K08HL153798 (P.C.), K08HL153798-S1(HuBMAP)(P.C.), R01HL171045 (T.Q), R01HL134817 (T.Q.), R01HL139478 (T.Q)., R01HL156846 (T.Q.), R01HL151535 (T.Q.), R01HL158525 (T.Q.), UM1 HG011972 (T.Q.), U01HG011762 (T.Q.), R01HL151535 (J.B.K), F32HL165854 (B.T.P), F32HL165819 (D.Y.L.); American Heart Association 23SCISA1144703(P.C), 24SCEFIA1248386 (P.C.), 20CDA35310303 (P.C.), 24CDA1272805 (B.T.P.); the Tobacco-Related Disease Research Program T32IR5352 (J.B.K), T32IR5240 (J.B.K);

## AUTHOR CONTRIBUTIONS

P.C., T.Q., and A.K conceived and supervised the study. Q.Z., D.S., and P.C performed bioinformatic analyses. P.C., A.P., W.G., A.D., C.W., W.J., A.B. and R.W. performed single cell captures, spatial transcriptomics, histology, and IHC. D.Y.L., H.S., and R.W. helped analyses. A.P., A.D., R.S. L.L. M.F. performed human tissue collection, D.Y.L., Y.R., T.N., R.S., B.T.P., J.P.M., M.W., A.B., M.I., R.K., L.L., helped conducting experiments. P.C., T.Q., and Q.Z. interpreted data and wrote the manuscript. R.W., M.F., and J.B.K contributed to experiments and manuscript. The authors read and approved the final manuscript.

## SUPPLEMENTAL INFORMATION

Figures S1–S10, Word file containing additional figure.

Figure S1. Canonical marker expression, donor variation, and spatial mapping of human arterial single-cell transcriptomes, related to Figure 1.

Figure S2. Smooth muscle cell (SMC) lineage heterogeneity, and characterization of microvasculature and SMC subtypes, related to Figure 2.

Figure S3. Fibroblast heterogeneity, subclustering, and functional divergence across vascular segments, related to Figure 3.

Figure S4. Characterization of endothelial cell identity, regional heterogeneity, lineage features, and disease associations, related to Figure 4.

Figure S5. Immune cell distribution and intercellular communication across human arterial segments.

Figure S6. Figure S6. Determinants of arterial identity across vascular territories.

Figure S7. The roles of embryonic origin in shaping transcriptomic diversity across arterial cell types, related to Figure 5.

Figure S8. Segment and cell type-specific patterns of vascular HOX gene expressions, related to Figure 5.

Figure S9. Cell type-specific enrichment of aortic aneurysm-associated and GWAS signals, related to Figure 6.

Figure S10. lncRNA expression patterns and their genetic associations with vascular disease, related to Figure 7.

Table S1-S12. Excel files containing additional data too large to fit in a PDF.

Table S1. Cell counts and proportions for each cluster across patients, related to Figure 1. Table S2. Differentially expressed genes between the two major clusters of SMC populations and their sub-clusters, related to Figure 2.

Table S3. Specific markers defined adventitial fibroblast population, related to Figure 3.

Table S4. Differentially expressed genes between the two endothelial cell populations, related to Figure 4.

Table S5. Motifs that are enriched in the promoters of genes differentially expressed between ascending and descending SMC or fibroblast.

Table S6. Differentially expressed genes between ascending and descending thoracic aorta, related to Figure 6.

Table S7. Diameter sensitive genes, related to Figure 6.

Table S8. Differentially expressed CAD GWAS genes, related to Figure 6. Table S9. Sex-specific differentially expressed genes.

Table S10. GO biological processes of sex-specific differentially expressed genes. Table S11. OMIM disease associations with sex-specific differentially expressed genes.

Table S12. lncRNAs overlapped with MVP GWAS loci for coronary artery diseases, related to Figure 7.

Table S13. The metadata for donors includes details about the collected tissues and the assays performed.

## STAR*METHODS

Detailed methods are provided in the online version of this paper and include the following:

- **KEY RESOURCES TABLE**
- **RESOURCE AVAILABILITY**

- Lead contact
- Materials availability
- Data and code availability
- **SUBJECT DETAILS**

- Human subjects
- **METHOD DETAILS**

- Human Tissue Collection
- Histology
- Single-Cell RNA Sequencing
- Analysis of Single-Cell Sequencing Data
- Spatial Slide-Seq
- Analysis of Spatial Slide-Seq Data
- Functional and Motif Analyses
- Correlation Analyses
- GWAS Enrichment Analyses
- Cell-cell Communications
- Lnc-RNA Analyses
- **QUANTIFICATION AND STATISTICAL ANALYSIS**

## REFERENCES

1. Tsao, C.W., Aday, A.W., Almarzooq, Z.I., Anderson, C.A.M., Arora, P., Avery, C.L., Baker-Smith, C.M., Beaton, A.Z., Boehme, A.K., Buxton, A.E., et al. (2023). Heart Disease and Stroke Statistics-2023 Update: A Report From the American Heart Association. Circulation 147, e93–e621. 10.1161/CIR.0000000000001123.

2. Cooke, J.P., and Meng, S. (2020). Vascular Regeneration in Peripheral Artery Disease. Arterioscler Thromb Vasc Biol 40, 1627–1634. 10.1161/ATVBAHA.120.312862.

3. De Backer, J., Bondue, A., Budts, W., Evangelista, A., Gallego, P., Jondeau, G., Loeys, B., Pena, M.L., Teixido-Tura, G., van de Laar, I., et al. (2020). Genetic counselling and testing in adults with congenital heart disease: A consensus document of the ESC Working Group of Grown-Up Congenital Heart Disease, the ESC Working Group on Aorta and Peripheral Vascular Disease and the European Society of Human Genetics. Eur J Prev Cardiol 27, 1423–1435. 10.1177/2047487319854552.

4. Donato, A.J., Machin, D.R., and Lesniewski, L.A. (2018). Mechanisms of Dysfunction in the Aging Vasculature and Role in Age-Related Disease. Circ Res 123, 825–848. 10.1161/CIRCRESAHA.118.312563.

5. Poredos, P., Poredos, P., and Jezovnik, M.K. (2018). Structure of Atherosclerotic Plaques in Different Vascular Territories: Clinical Relevance. Curr Vasc Pharmacol 16, 125–129. 10.2174/1570161115666170227103125.

6. DeBakey, M.E., Lawrie, G.M., and Glaeser, D.H. (1985). Patterns of atherosclerosis and their surgical significance. Ann Surg 201, 115–131. 10.1097/00000658-198502000-00001.

7. Morisaki, T., and Morisaki, H. (2016). Genetics of hereditary large vessel diseases. J Hum Genet 61, 21–26. 10.1038/jhg.2015.119.

8. Loeys, B.L., Schwarze, U., Holm, T., Callewaert, B.L., Thomas, G.H., Pannu, H., De Backer, J.F., Oswald, G.L., Symoens, S., Manouvrier, S., et al. (2006). Aneurysm syndromes caused by mutations in the TGF-beta receptor. N Engl J Med 355, 788–798. 10.1056/NEJMoa055695.

9. Kim, J.B., Zhao, Q., Nguyen, T., Pjanic, M., Cheng, P., Wirka, R., Travisano, S., Nagao, M., Kundu, R., and Quertermous, T. (2020). Environment-Sensing Aryl Hydrocarbon Receptor Inhibits the Chondrogenic Fate of Modulated Smooth Muscle Cells in Atherosclerotic Lesions. Circulation 142, 575–590. 10.1161/CIRCULATIONAHA.120.045981.

10. Kondo, T., Nakano, Y., Adachi, S., and Murohara, T. (2019). Effects of Tobacco Smoking on Cardiovascular Disease. Circ J 83, 1980–1985. 10.1253/circj.CJ-19-0323.

11. Maleszewski, J.J. (2015). Inflammatory ascending aortic disease: perspectives from pathology. J Thorac Cardiovasc Surg 149, S176–183. 10.1016/j.jtcvs.2014.07.046.

12. Rajasekaran, K., Duraiyarasan, S., Adefuye, M., Manjunatha, N., and Ganduri, V. (2022). Kawasaki Disease and Coronary Artery Involvement: A Narrative Review. Cureus 14, e28358. 10.7759/cureus.28358.

13. Yuan, S.M. (2018). Syphilitic aortic aneurysm. Z Rheumatol 77, 741–748. 10.1007/s00393-018-0519-1.

14. Noden, D.M. (1989). Embryonic origins and assembly of blood vessels. Am Rev Respir Dis 140, 1097–1103. 10.1164/ajrccm/140.4.1097.

15. Saha, M.S., Cox, E.A., and Sipe, C.W. (2004). Mechanisms regulating the origins of the vertebrate vascular system. J Cell Biochem 93, 46–56. 10.1002/jcb.20196.

16. Leonard, E.V., Figueroa, R.J., Bussmann, J., Lawson, N.D., Amigo, J.D., and Siekmann, A.F. (2022). Regenerating vascular mural cells in zebrafish fin blood vessels are not derived from pre-existing mural cells and differentially require Pdgfrb signalling for their development. Development 149. 10.1242/dev.199640.

17. Miano, J.M., Fisher, E.A., and Majesky, M.W. (2021). Fate and State of Vascular Smooth Muscle Cells in Atherosclerosis. Circulation 143, 2110–2116. 10.1161/CIRCULATIONAHA.120.049922.

18. Rasanen, M., Sultan, I., Paech, J., Hemanthakumar, K.A., Yu, W., He, L., Tang, J., Sun, Y., Hlushchuk, R., Huan, X., et al. (2021). VEGF-B Promotes Endocardium-Derived Coronary Vessel Development and Cardiac Regeneration. Circulation 143, 65–77. 10.1161/CIRCULATIONAHA.120.050635.

19. Sharma, B., Chang, A., and Red-Horse, K. (2017). Coronary Artery Development: Progenitor Cells and Differentiation Pathways. Annu Rev Physiol 79, 1–19. 10.1146/annurev-physiol-022516-033953.

20. Red-Horse, K., Ueno, H., Weissman, I.L., and Krasnow, M.A. (2010). Coronary arteries form by developmental reprogramming of venous cells. Nature 464, 549–553. 10.1038/nature08873.

21. Majesky, M.W. (2007). Developmental basis of vascular smooth muscle diversity. Arterioscler Thromb Vasc Biol 27, 1248–1258. 10.1161/ATVBAHA.107.141069.

22. High, F.A., Lu, M.M., Pear, W.S., Loomes, K.M., Kaestner, K.H., and Epstein, J.A. (2008). Endothelial expression of the Notch ligand Jagged1 is required for vascular smooth muscle development. Proc Natl Acad Sci U S A 105, 1955–1959. 10.1073/pnas.0709663105.

23. Sato, Y. (2013). Dorsal aorta formation: separate origins, lateral-to-medial migration, and remodeling. Dev Growth Differ 55, 113–129. 10.1111/dgd.12010.

24. Dobnikar, L., Taylor, A.L., Chappell, J., Oldach, P., Harman, J.L., Oerton, E., Dzierzak, E., Bennett, M.R., Spivakov, M., and Jorgensen, H.F. (2018). Disease-relevant transcriptional signatures identified in individual smooth muscle cells from healthy mouse vessels. Nat Commun 9, 4567. 10.1038/s41467-018-06891-x.

25. Trigueros-Motos, L., Gonzalez-Granado, J.M., Cheung, C., Fernandez, P., Sanchez- Cabo, F., Dopazo, A., Sinha, S., and Andres, V. (2013). Embryological-origin-dependent differences in homeobox expression in adult aorta: role in regional phenotypic variability and regulation of NF-kappaB activity. Arterioscler Thromb Vasc Biol 33, 1248–1256. 10.1161/ATVBAHA.112.300539.

26. Haimovici, H., Maier, N., and Strauss, L. (1959). Fate of aortic homografts in experimental canine atherosclerosis. II. Study of fresh abdominal aortic implants into abdominal aorta. AMA Arch Surg 78, 239–245. 10.1001/archsurg.1959.04320020061010.

27. Haimovici, H., and Maier, N. (1964). Fate of Aortic Homografts in Canine Atherosclerosis. 3. Study of Fresh Abdominal and Thoracic Aortic Implants into Thoracic Aorta: Role of Tissue Susceptibility in Atherogenesis. Arch Surg 89, 961–969. 10.1001/archsurg.1964.01320060029006.

28. Wirka, R.C., Wagh, D., Paik, D.T., Pjanic, M., Nguyen, T., Miller, C.L., Kundu, R., Nagao, M., Coller, J., Koyano, T.K., et al. (2019). Atheroprotective roles of smooth muscle cell phenotypic modulation and the TCF21 disease gene as revealed by single-cell analysis. Nat Med 25, 1280–1289. 10.1038/s41591-019-0512-5.

29. Cheng, P., Wirka, R.C., Kim, J.B., Kim, H.J., Nguyen, T., Kundu, R., Zhao, Q., Sharma, D., Pedroza, A., Nagao, M., et al. (2022). Smad3 regulates smooth muscle cell fate and mediates adverse remodeling and calcification of the atherosclerotic plaque. Nat Cardiovasc Res 1, 322–333. 10.1038/s44161-022-00042-8.

30. Cheng, P., Wirka, R.C., Shoa Clarke, L., Zhao, Q., Kundu, R., Nguyen, T., Nair, S., Sharma, D., Kim, H.J., Shi, H., et al. (2022). ZEB2 Shapes the Epigenetic Landscape of Atherosclerosis. Circulation 145, 469–485. 10.1161/CIRCULATIONAHA.121.057789.

31. Hafemeister, C., and Satija, R. (2019). Normalization and variance stabilization of single- cell RNA-seq data using regularized negative binomial regression. Genome Biol 20, 296. 10.1186/s13059-019-1874-1.

32. Soliman, H., Tung, L.W., and Rossi, F.M.V. (2021). Fibroblast and Myofibroblast Subtypes: Single Cell Sequencing. Methods Mol Biol 2299, 49–84. 10.1007/978-1-0716-1382-5_4.

33. Hume, D.A., Millard, S.M., and Pettit, A.R. (2023). Macrophage heterogeneity in the single-cell era: facts and artifacts. Blood 142, 1339–1347. 10.1182/blood.2023020597.

34. Mae, M.A., He, L., Nordling, S., Vazquez-Liebanas, E., Nahar, K., Jung, B., Li, X., Tan, B.C., Chin Foo, J., Cazenave-Gassiot, A., et al. (2021). Single-Cell Analysis of Blood- Brain Barrier Response to Pericyte Loss. Circ Res 128, e46–e62. 10.1161/CIRCRESAHA.120.317473.

35. Kalucka, J., de Rooij, L., Goveia, J., Rohlenova, K., Dumas, S.J., Meta, E., Conchinha, N.V., Taverna, F., Teuwen, L.A., Veys, K., et al. (2020). Single-Cell Transcriptome Atlas of Murine Endothelial Cells. Cell 180, 764–779 e720. 10.1016/j.cell.2020.01.015.

36. Muhl, L., Mocci, G., Pietila, R., Liu, J., He, L., Genove, G., Leptidis, S., Gustafsson, S., Buyandelger, B., Raschperger, E., et al. (2022). A single-cell transcriptomic inventory of murine smooth muscle cells. Dev Cell 57, 2426–2443 e2426. 10.1016/j.devcel.2022.09.015.

37. Allahverdian, S., Chaabane, C., Boukais, K., Francis, G.A., and Bochaton-Piallat, M.L. (2018). Smooth muscle cell fate and plasticity in atherosclerosis. Cardiovasc Res 114, 540–550. 10.1093/cvr/cvy022.

38. Hu, Z., Liu, W., Hua, X., Chen, X., Chang, Y., Hu, Y., Xu, Z., and Song, J. (2021). Single-Cell Transcriptomic Atlas of Different Human Cardiac Arteries Identifies Cell Types Associated With Vascular Physiology. Arterioscler Thromb Vasc Biol 41, 1408–1427. 10.1161/ATVBAHA.120.315373.

39. Crnkovic, S., Valzano, F., Fliesser, E., Gindlhuber, J., Thekkekara Puthenparampil, H., Basil, M., Morley, M.P., Katzen, J., Gschwandtner, E., Klepetko, W., et al. (2022). Single-cell transcriptomics reveals skewed cellular communication and phenotypic shift in pulmonary artery remodeling. JCI Insight 7. 10.1172/jci.insight.153471.

40. Chakraborty, R., Saddouk, F.Z., Carrao, A.C., Krause, D.S., Greif, D.M., and Martin, K.A. (2019). Promoters to Study Vascular Smooth Muscle. Arterioscler Thromb Vasc Biol 39, 603–612. 10.1161/ATVBAHA.119.312449.

41. Armulik, A., Genove, G., and Betsholtz, C. (2011). Pericytes: developmental, physiological, and pathological perspectives, problems, and promises. Dev Cell 21, 193–215. 10.1016/j.devcel.2011.07.001.

42. Shankman, L.S., Gomez, D., Cherepanova, O.A., Salmon, M., Alencar, G.F., Haskins, R.M., Swiatlowska, P., Newman, A.A., Greene, E.S., Straub, A.C., et al. (2015). KLF4- dependent phenotypic modulation of smooth muscle cells has a key role in atherosclerotic plaque pathogenesis. Nat Med 21, 628–637. 10.1038/nm.3866.

43. Kramann, R., Schneider, R.K., DiRocco, D.P., Machado, F., Fleig, S., Bondzie, P.A., Henderson, J.M., Ebert, B.L., and Humphreys, B.D. (2015). Perivascular Gli1+ progenitors are key contributors to injury-induced organ fibrosis. Cell Stem Cell 16, 51–66. 10.1016/j.stem.2014.11.004.

44. Guerrero-Juarez, C.F., Dedhia, P.H., Jin, S., Ruiz-Vega, R., Ma, D., Liu, Y., Yamaga, K., Shestova, O., Gay, D.L., Yang, Z., et al. (2019). Single-cell analysis reveals fibroblast heterogeneity and myeloid-derived adipocyte progenitors in murine skin wounds. Nat Commun 10, 650. 10.1038/s41467-018-08247-x.

45. Xie, T., Wang, Y., Deng, N., Huang, G., Taghavifar, F., Geng, Y., Liu, N., Kulur, V., Yao, C., Chen, P., et al. (2018). Single-Cell Deconvolution of Fibroblast Heterogeneity in Mouse Pulmonary Fibrosis. Cell Rep 22, 3625–3640. 10.1016/j.celrep.2018.03.010.

46. Philippeos, C., Telerman, S.B., Oules, B., Pisco, A.O., Shaw, T.J., Elgueta, R., Lombardi, G., Driskell, R.R., Soldin, M., Lynch, M.D., and Watt, F.M. (2018). Spatial and Single- Cell Transcriptional Profiling Identifies Functionally Distinct Human Dermal Fibroblast Subpopulations. J Invest Dermatol 138, 811–825. 10.1016/j.jid.2018.01.016.

47. Pedroza, A.J., Tashima, Y., Shad, R., Cheng, P., Wirka, R., Churovich, S., Nakamura, K., Yokoyama, N., Cui, J.Z., Iosef, C., et al. (2020). Single-Cell Transcriptomic Profiling of Vascular Smooth Muscle Cell Phenotype Modulation in Marfan Syndrome Aortic Aneurysm. Arterioscler Thromb Vasc Biol 40, 2195–2211. 10.1161/ATVBAHA.120.314670.

48. Renner, S., Schuler, H., Alawi, M., Kolbe, V., Rybczynski, M., Woitschach, R., Sheikhzadeh, S., Stark, V.C., Olfe, J., Roser, E., et al. (2019). Next-generation sequencing of 32 genes associated with hereditary aortopathies and related disorders of connective tissue in a cohort of 199 patients. Genet Med 21, 1832–1841. 10.1038/s41436-019-0435-z.

49. Sawada, H., Katsumata, Y., Higashi, H., Zhang, C., Li, Y., Morgan, S., Lee, L.H., Singh, S.A., Chen, J.Z., Franklin, M.K., et al. (2022). Second Heart Field-Derived Cells Contribute to Angiotensin II-Mediated Ascending Aortopathies. Circulation 145, 987–1001. 10.1161/CIRCULATIONAHA.121.058173.

50. Naito, H., Iba, T., and Takakura, N. (2020). Mechanisms of new blood-vessel formation and proliferative heterogeneity of endothelial cells. Int Immunol 32, 295–305. 10.1093/intimm/dxaa008.

51. Hennigs, J.K., Matuszcak, C., Trepel, M., and Korbelin, J. (2021). Vascular Endothelial Cells: Heterogeneity and Targeting Approaches. Cells 10. 10.3390/cells10102712.

52. Fischer, A., Schumacher, N., Maier, M., Sendtner, M., and Gessler, M. (2004). The Notch target genes Hey1 and Hey2 are required for embryonic vascular development. Genes Dev 18, 901–911. 10.1101/gad.291004.

53. Dong, G., Huang, X., Xu, Y., Chen, R., and Chen, S. (2023). Mechanical stress induced EndoMT in endothelial cells through PPARgamma downregulation. Cell Signal 110, 110812. 10.1016/j.cellsig.2023.110812.

54. Li, Y., Lui, K.O., and Zhou, B. (2018). Reassessing endothelial-to-mesenchymal transition in cardiovascular diseases. Nat Rev Cardiol 15, 445–456. 10.1038/s41569-018-0023-y.

55. Xi, N.M., and Li, J.J. (2021). Benchmarking Computational Doublet-Detection Methods for Single-Cell RNA Sequencing Data. Cell Syst 12, 176–194 e176. 10.1016/j.cels.2020.11.008.

56. Arroz-Madeira, S., Bekkhus, T., Ulvmar, M.H., and Petrova, T.V. (2023). Lessons of Vascular Specialization From Secondary Lymphoid Organ Lymphatic Endothelial Cells. Circ Res 132, 1203–1225. 10.1161/CIRCRESAHA.123.322136.

57. Jalkanen, S., and Salmi, M. (2020). Lymphatic endothelial cells of the lymph node. Nat Rev Immunol 20, 566–578. 10.1038/s41577-020-0281-x.

58. Hedhli, N., Kalinowski, A., and K, S.R. (2014). Cardiovascular effects of neuregulin- 1/ErbB signaling: role in vascular signaling and angiogenesis. Curr Pharm Des 20, 4899–4905. 10.2174/1381612819666131125151058.

59. Odiete, O., Hill, M.F., and Sawyer, D.B. (2012). Neuregulin in cardiovascular development and disease. Circ Res 111, 1376–1385. 10.1161/CIRCRESAHA.112.267286.

60. Guclu-Geyik, F., Erkan, A.F., Ozuynuk, A.S., Ekici, B., and Coban, N. (2022). Val109Asp polymorphism in Intelectin 1 gene is associated with coronary artery disease severity in women. Turk Kardiyol Dern Ars 50, 34–45. 10.5543/tkda.2022.21003.

61. Gu, N., Dong, Y., Tian, Y., Di, Z., Liu, Z., Chang, M., Jia, X., Qian, Y., and Zhang, W. (2017). Anti-apoptotic and angiogenic effects of intelectin-1 in rat cerebral ischemia. Brain Res Bull 130, 27–35. 10.1016/j.brainresbull.2016.12.006.

62. Luo, X.L., Jiang, Y., Li, Q., Yu, X.J., Ma, T., Cao, H., Ke, M.X., Zhang, P., Tan, J.L., Gong, Y.S., et al. (2023). hESC-Derived Epicardial Cells Promote Repair of Infarcted Hearts in Mouse and Swine. Adv Sci (Weinh) 10, e2300470. 10.1002/advs.202300470.

63. Menzel, J., di Giuseppe, R., Biemann, R., Wittenbecher, C., Aleksandrova, K., Pischon, T., Fritsche, A., Schulze, M.B., Boeing, H., Isermann, B., and Weikert, C. (2016). Omentin-1 and risk of myocardial infarction and stroke: Results from the EPIC-Potsdam cohort study. Atherosclerosis 251, 415–421. 10.1016/j.atherosclerosis.2016.06.003.

64. Litvinukova, M., Talavera-Lopez, C., Maatz, H., Reichart, D., Worth, C.L., Lindberg, E.L., Kanda, M., Polanski, K., Heinig, M., Lee, M., et al. (2020). Cells of the adult human heart. Nature 588, 466–472. 10.1038/s41586-020-2797-4.

65. Zhang, X., McDonald, J.G., Aryal, B., Canfran-Duque, A., Goldberg, E.L., Araldi, E., Ding, W., Fan, Y., Thompson, B.M., Singh, A.K., et al. (2021). Desmosterol suppresses macrophage inflammasome activation and protects against vascular inflammation and atherosclerosis. Proc Natl Acad Sci U S A 118. 10.1073/pnas.2107682118.

66. Zhang, Q., Liu, Q., Feng, S., Li, X., and Jiang, Q. (2024). Non-Coding RNAs: Novel Regulators of Macrophage Homeostasis in Ocular Vascular Diseases. Biomolecules 14. 10.3390/biom14030328.

67. Jin, S., Guerrero-Juarez, C.F., Zhang, L., Chang, I., Ramos, R., Kuan, C.H., Myung, P., Plikus, M.V., and Nie, Q. (2021). Inference and analysis of cell-cell communication using CellChat. Nat Commun 12, 1088. 10.1038/s41467-021-21246-9.

68. Becht, E., McInnes, L., Healy, J., Dutertre, C.A., Kwok, I.W.H., Ng, L.G., Ginhoux, F., and Newell, E.W. (2018). Dimensionality reduction for visualizing single-cell data using UMAP. Nat Biotechnol. 10.1038/nbt.4314.

69. Zhai, Z., Lei, Y.L., Wang, R., and Xie, Y. (2022). Supervised capacity preserving mapping: a clustering guided visualization method for scRNA-seq data. Bioinformatics 38, 2496–2503. 10.1093/bioinformatics/btac131.

70. Liu, J., and Vinck, M. (2022). Improved visualization of high-dimensional data using the distance-of-distance transformation. PLoS Comput Biol 18, e1010764. 10.1371/journal.pcbi.1010764.

71. Ultsch, A., and Lotsch, J. (2022). Euclidean distance-optimized data transformation for cluster analysis in biomedical data (EDOtrans). BMC Bioinformatics 23, 233. 10.1186/s12859-022-04769-w.

72. Gorski, D.H., and Walsh, K. (2000). The role of homeobox genes in vascular remodeling and angiogenesis. Circ Res 87, 865–872. 10.1161/01.res.87.10.865.

73. Swift, M.R., and Weinstein, B.M. (2009). Arterial-venous specification during development. Circ Res 104, 576–588. 10.1161/CIRCRESAHA.108.188805.

74. Barnes, R.M., Firulli, B.A., VanDusen, N.J., Morikawa, Y., Conway, S.J., Cserjesi, P., Vincentz, J.W., and Firulli, A.B. (2011). Hand2 loss-of-function in Hand1-expressing cells reveals distinct roles in epicardial and coronary vessel development. Circ Res 108, 940–949. 10.1161/CIRCRESAHA.110.233171.

75. Maruyama, K., Miyagawa-Tomita, S., Haneda, Y., Kida, M., Matsuzaki, F., Imanaka- Yoshida, K., and Kurihara, H. (2022). The cardiopharyngeal mesoderm contributes to lymphatic vessel development in mouse. Elife 11. 10.7554/eLife.81515.

76. Materna, S.C., Sinha, T., Barnes, R.M., Lammerts van Bueren, K., and Black, B.L. (2019). Cardiovascular development and survival require Mef2c function in the myocardial but not the endothelial lineage. Dev Biol 445, 170–177. 10.1016/j.ydbio.2018.12.002.

77. Crispino, J.D., Lodish, M.B., Thurberg, B.L., Litovsky, S.H., Collins, T., Molkentin, J.D., and Orkin, S.H. (2001). Proper coronary vascular development and heart morphogenesis depend on interaction of GATA-4 with FOG cofactors. Genes Dev 15, 839–844. 10.1101/gad.875201.

78. Hubert, K.A., and Wellik, D.M. (2023). Hox genes in development and beyond. Development 150. 10.1242/dev.192476.

79. Vapnik, J.S., Kim, J.B., Isselbacher, E.M., Ghoshhajra, B.B., Cheng, Y., Sundt, T.M., 3rd, MacGillivray, T.E., Cambria, R.P., and Lindsay, M.E. (2016). Characteristics and Outcomes of Ascending Versus Descending Thoracic Aortic Aneurysms. Am J Cardiol 117, 1683–1690. 10.1016/j.amjcard.2016.02.048.

80. Huang, Y., Schaff, H.V., Bagameri, G., Pochettino, A., DeMartino, R.R., Todd, A., and Greason, K.L. (2024). Differential expansion and outcomes of ascending and descending degenerative thoracic aortic aneurysms. J Thorac Cardiovasc Surg 167, 918–926 e913. 10.1016/j.jtcvs.2022.03.032.

81. Chou, E.L., and Lindsay, M.E. (2020). The genetics of aortopathies: Hereditary thoracic aortic aneurysms and dissections. Am J Med Genet C Semin Med Genet 184, 136–148. 10.1002/ajmg.c.31771.

82. Isselbacher, E.M., Lino Cardenas, C.L., and Lindsay, M.E. (2016). Hereditary Influence in Thoracic Aortic Aneurysm and Dissection. Circulation 133, 2516–2528. 10.1161/CIRCULATIONAHA.116.009762.

83. Shalhub, S., Regalado, E.S., Guo, D.C., Milewicz, D.M., and Montalcino Aortic, C. (2019). The natural history of type B aortic dissection in patients with PRKG1 mutation c.530G>A (p.Arg177Gln). J Vasc Surg 70, 718–723. 10.1016/j.jvs.2018.12.032.

84. Gago-Diaz, M., Blanco-Verea, A., Teixido, G., Huguet, F., Gut, M., Laurie, S., Gut, I., Carracedo, A., Evangelista, A., and Brion, M. (2016). PRKG1 and genetic diagnosis of early-onset thoracic aortic disease. Eur J Clin Invest 46, 787–794. 10.1111/eci.12662.

85. Pirruccello, J.P., Chaffin, M.D., Chou, E.L., Fleming, S.J., Lin, H., Nekoui, M., Khurshid, S., Friedman, S.F., Bick, A.G., Arduini, A., et al. (2022). Deep learning enables genetic analysis of the human thoracic aorta. Nat Genet 54, 40–51. 10.1038/s41588-021-00962-4.

86. Khoury, Z., Gottlieb, S., Stern, S., and Keren, A. (1997). Frequency and distribution of atherosclerotic plaques in the thoracic aorta as determined by transesophageal echocardiography in patients with coronary artery disease. Am J Cardiol 79, 23–27. 10.1016/s0002-9149(96)00670-4.

87. Allais, G., Chiarle, G., Sinigaglia, S., Airola, G., Schiapparelli, P., and Benedetto, C. (2018). Estrogen, migraine, and vascular risk. Neurol Sci 39, 11–20. 10.1007/s10072-018-3333-2.

88. Klouche, M. (2006). Estrogens in human vascular diseases. Ann N Y Acad Sci 1089, 431–443. 10.1196/annals.1386.032.

89. Lin, G.M., Yano, Y., and Hoshide, S. (2017). Sex Differences in the Association between Traditional Vascular Risk Factors and Subclinical Carotid Atherosclerosis in Taiwan. J Atheroscler Thromb 24, 673–674. 10.5551/jat.ED068.

90. Liu, P.P., and Fukuoka, M. (2007). Sex hormones as novel risk biomarkers for atherosclerosis in peripheral vascular disease. J Am Coll Cardiol 50, 1077–1079. 10.1016/j.jacc.2007.05.031.

91. Yahagi, K., Davis, H.R., Arbustini, E., and Virmani, R. (2015). Sex differences in coronary artery disease: pathological observations. Atherosclerosis 239, 260–267. 10.1016/j.atherosclerosis.2015.01.017.

92. Nucera, M., Heinisch, P.P., Langhammer, B., Jungi, S., Mihalj, M., Schober, P., Luedi, M.M., Yildiz, M., and Schoenhoff, F.S. (2022). The impact of sex and gender on aortic events in patients with Marfan syndrome. Eur J Cardiothorac Surg 62. 10.1093/ejcts/ezac305.

93. Shimizu, Y., and Murohara, T. (2024). Takayasu Arteritis in Terms of Disease Duration and Sex Differences. Circ J 88, 295–296. 10.1253/circj.CJ-23-0900.

94. Brundage, J.N., Mason, C.P., Wadsworth, H.A., Finuf, C.S., Nelson, J.J., Ronstrom, P.J.W., Jones, S.R., Siciliano, C.A., Steffensen, S.C., and Yorgason, J.T. (2022). Regional and sex differences in spontaneous striatal dopamine transmission. J Neurochem 160, 598–612. 10.1111/jnc.15473.

95. Crouse, J.R., 3rd, Craven, T.E., Hagaman, A.P., and Bond, M.G. (1995). Association of coronary disease with segment-specific intimal-medial thickening of the extracranial carotid artery. Circulation 92, 1141–1147. 10.1161/01.cir.92.5.1141.

96. Nakashima, Y., Wight, T.N., and Sueishi, K. (2008). Early atherosclerosis in humans: role of diffuse intimal thickening and extracellular matrix proteoglycans. Cardiovasc Res 79, 14–23. 10.1093/cvr/cvn099.

97. Milutinovic, A., Suput, D., and Zorc-Pleskovic, R. (2020). Pathogenesis of atherosclerosis in the tunica intima, media, and adventitia of coronary arteries: An updated review. Bosn J Basic Med Sci 20, 21–30. 10.17305/bjbms.2019.4320.

98. Shi, H., Nguyen, T., Zhao, Q., Cheng, P., Sharma, D., Kim, H.J., Brian Kim, J., Wirka, R., Weldy, C.S., Monteiro, J.P., and Quertermous, T. (2023). Discovery of Transacting Long Noncoding RNAs That Regulate Smooth Muscle Cell Phenotype. Circ Res 132, 795–811. 10.1161/CIRCRESAHA.122.321960.

99. Leung, A., and Natarajan, R. (2014). Noncoding RNAs in vascular disease. Curr Opin Cardiol 29, 199–206. 10.1097/HCO.0000000000000054.

100. Simion, V., Haemmig, S., and Feinberg, M.W. (2019). LncRNAs in vascular biology and disease. Vascul Pharmacol 114, 145–156. 10.1016/j.vph.2018.01.003.

101. Sollis, E., Mosaku, A., Abid, A., Buniello, A., Cerezo, M., Gil, L., Groza, T., Gunes, O., Hall, P., Hayhurst, J., et al. (2023). The NHGRI-EBI GWAS Catalog: knowledgebase and deposition resource. Nucleic Acids Res 51, D977–D985. 10.1093/nar/gkac1010.

102. Djousse, L., Ho, Y.L., Nguyen, X.T., Gagnon, D.R., Wilson, P.W.F., Cho, K., Gaziano, J.M., and Program, V.A.M.V. (2018). DASH Score and Subsequent Risk of Coronary Artery Disease: The Findings From Million Veteran Program. J Am Heart Assoc 7. 10.1161/JAHA.117.008089.

103. Nikpay, M., Goel, A., Won, H.H., Hall, L.M., Willenborg, C., Kanoni, S., Saleheen, D., Kyriakou, T., Nelson, C.P., Hopewell, J.C., et al. (2015). A comprehensive 1,000 Genomes-based genome-wide association meta-analysis of coronary artery disease. Nat Genet 47, 1121–1130. 10.1038/ng.3396.

104. Petkovic, A., Erceg, S., Munjas, J., Ninic, A., Vladimirov, S., Davidovic, A., Vukmirovic, L., Milanov, M., Cvijanovic, D., Mitic, T., and Sopic, M. (2023). LncRNAs as Regulators of Atherosclerotic Plaque Stability. Cells 12. 10.3390/cells12141832.

105. Ehler, E., Babiychuk, E., and Draeger, A. (1996). Human foetal lung (IMR-90) cells: myofibroblasts with smooth muscle-like contractile properties. Cell Motil Cytoskeleton 34, 288–298. 10.1002/(SICI)1097-0169(1996)34:4<288::AID-CM4>3.0.CO;2-4.

106. Roelofs, M., Faggian, L., Pampinella, F., Paulon, T., Franch, R., Chiavegato, A., and Sartore, S. (1998). Transforming growth factor beta1 involvement in the conversion of fibroblasts to smooth muscle cells in the rabbit bladder serosa. Histochem J 30, 393–404. 10.1023/a:1003216124761.

107. Varberg, K.M., Garretson, R.O., Blue, E.K., Chu, C., Gohn, C.R., Tu, W., and Haneline, L.S. (2018). Transgelin induces dysfunction of fetal endothelial colony-forming cells from gestational diabetic pregnancies. Am J Physiol Cell Physiol 315, C502–C515. 10.1152/ajpcell.00137.2018.

108. Warthi, G., Faulkner, J.L., Doja, J., Ghanam, A.R., Gao, P., Yang, A.C., Slivano, O.J., Barris, C.T., Kress, T.C., Zawieja, S.D., et al. (2022). Generation and Comparative Analysis of an Itga8-CreER (T2) Mouse with Preferential Activity in Vascular Smooth Muscle Cells. Nat Cardiovasc Res 1, 1084–1100. 10.1038/s44161-022-00162-1.

109. Avila Cobos, F., Alquicira-Hernandez, J., Powell, J.E., Mestdagh, P., and De Preter, K. (2020). Benchmarking of cell type deconvolution pipelines for transcriptomics data. Nat Commun 11, 5650. 10.1038/s41467-020-19015-1.

110. Wang, X., Park, J., Susztak, K., Zhang, N.R., and Li, M. (2019). Bulk tissue cell type deconvolution with multi-subject single-cell expression reference. Nat Commun 10, 380. 10.1038/s41467-018-08023-x.

111. Gaudio, L.T., Caruso, M.V., De Rosa, S., Indolfi, C., and Fragomeni, G. (2018). Different Blood Flow Models in Coronary Artery Diseases: Effects on hemodynamic parameters. Annu Int Conf IEEE Eng Med Biol Soc 2018, 3185–3188. 10.1109/EMBC.2018.8512917.

112. Govindaraju, K., Badruddin, I.A., Viswanathan, G.N., Ramesh, S.V., and Badarudin, A. (2013). Evaluation of functional severity of coronary artery disease and fluid dynamics’ influence on hemodynamic parameters: A review. Phys Med 29, 225–232. 10.1016/j.ejmp.2012.03.008.

113. Nixon, A.M., Gunel, M., and Sumpio, B.E. (2010). The critical role of hemodynamics in the development of cerebral vascular disease. J Neurosurg 112, 1240–1253. 10.3171/2009.10.JNS09759.

114. Chiu, J.J., and Chien, S. (2011). Effects of disturbed flow on vascular endothelium: pathophysiological basis and clinical perspectives. Physiol Rev 91, 327–387. 10.1152/physrev.00047.2009.

115. Wang, C., Baker, B.M., Chen, C.S., and Schwartz, M.A. (2013). Endothelial cell sensing of flow direction. Arterioscler Thromb Vasc Biol 33, 2130–2136. 10.1161/ATVBAHA.113.301826.

116. Langley-Evans, S.C., and McMullen, S. (2010). Developmental origins of adult disease. Med Princ Pract 19, 87–98. 10.1159/000273066.

117. Thornburg, K.L. (2015). The programming of cardiovascular disease. J Dev Orig Health Dis 6, 366–376. 10.1017/S2040174415001300.

118. Tian, X., Pu, W.T., and Zhou, B. (2015). Cellular origin and developmental program of coronary angiogenesis. Circ Res 116, 515–530. 10.1161/CIRCRESAHA.116.305097.

119. MacFarlane, E.G., Parker, S.J., Shin, J.Y., Kang, B.E., Ziegler, S.G., Creamer, T.J., Bagirzadeh, R., Bedja, D., Chen, Y., Calderon, J.F., et al. (2019). Lineage-specific events underlie aortic root aneurysm pathogenesis in Loeys-Dietz syndrome. J Clin Invest 129, 659–675. 10.1172/JCI123547.

120. Gailloud, P. (2022). The segmentation of the vertebral artery: An ambiguous anatomical concept. Interv Neuroradiol 28, 765–772. 10.1177/15910199211063275.

121. Liu, M., and Gomez, D. (2019). Smooth Muscle Cell Phenotypic Diversity. Arterioscler Thromb Vasc Biol 39, 1715–1723. 10.1161/ATVBAHA.119.312131.

122. Bentzon, J.F., and Majesky, M.W. (2018). Lineage tracking of origin and fate of smooth muscle cells in atherosclerosis. Cardiovasc Res 114, 492–500. 10.1093/cvr/cvx251.

123. Egusa, G., Watanabe, H., Ohshita, K., Fujikawa, R., Yamane, K., Okubo, M., and Kohno, N. (2002). Influence of the extent of westernization of lifestyle on the progression of preclinical atherosclerosis in Japanese subjects. J Atheroscler Thromb 9, 299–304. 10.5551/jat.9.299.

