## Supplemental Figures for "A cell and transcriptome atlas of human arterial vasculature"

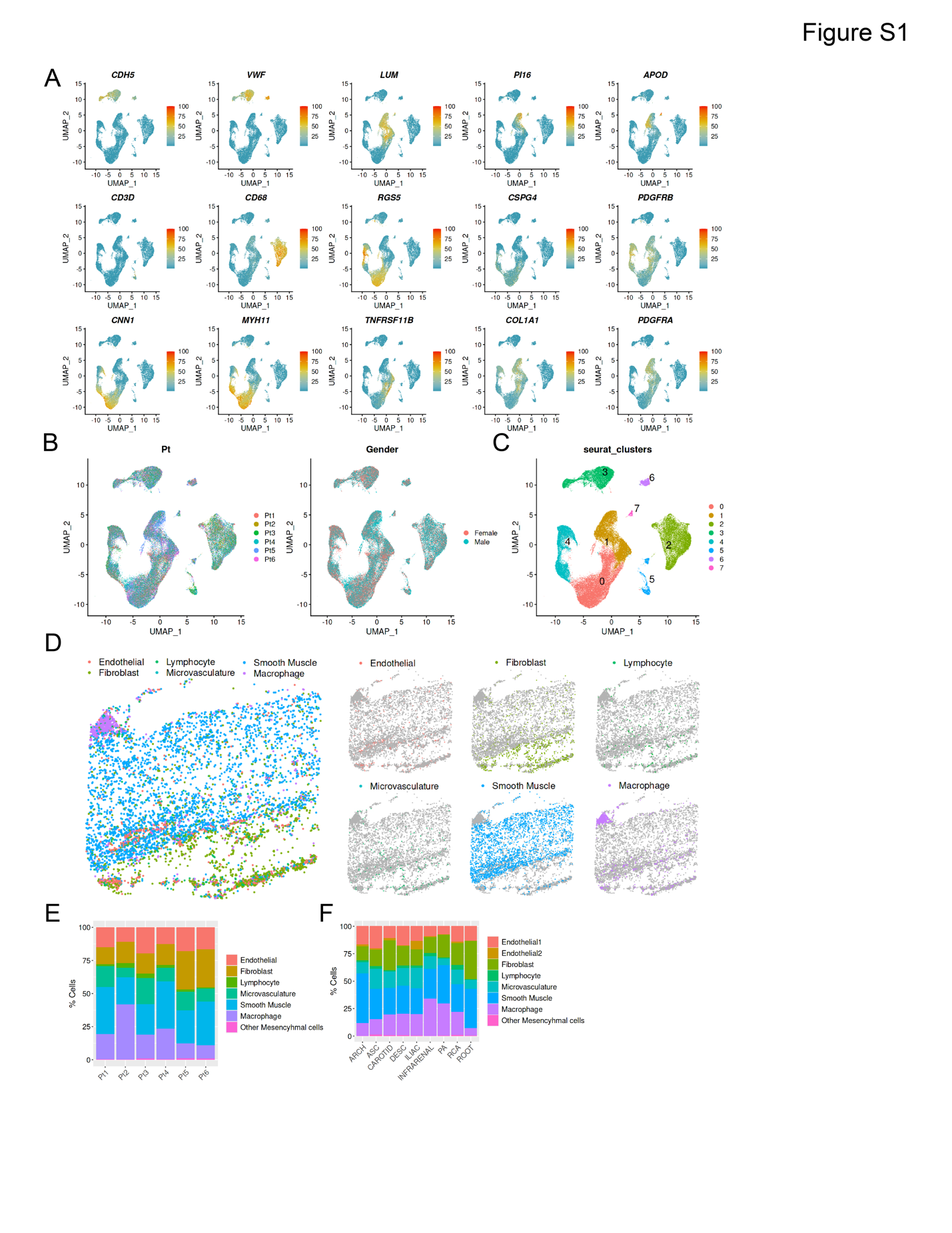

**Figure S1. Canonical marker expression, donor variation, and spatial mapping of human arterial single-cell transcriptomes.**

(A) UMAP visualization showing the expression patterns of canonical marker genes used to define each major vascular cell population, including smooth muscle cells (e.g., *MYH11*, *ACTA2*), fibroblasts (e.g., *FBLN1*, *LOX*), endothelial cells (e.g., *CDH5*, *VWF*), and immune cells (e.g., *PTPRC*, *CD68*).

(B) Uniform Manifold Approximation and Projection (UMAP) plots illustrating inter-individual heterogeneity. Left: clustering of cells colored by donor (Pt), showing subject-specific variation in transcriptomic profiles. Right: clustering colored by sex, suggesting gender-associated differences in cellular composition or gene expression.

(C) UMAP representation of the initial low-resolution clustering generated by Seurat prior to cell type annotation, demonstrating the underlying transcriptomic structure of the dataset.

(D) Spatial dim plot of a representative aortic section. Left panel: spatial annotation of each cell type. Right panel: split view showing the discrete spatial distribution of individual cell types, highlighting their anatomical niches within the tissue.

(E) Bar plot summarizing the proportion of each cell type in the six individual donors, revealing commonalities in cell composition.

(F) Bar plot showing the distribution of each major cell population across all vascular beds included in the study.

Abbreviations: ascending thoracic aorta (ASC), aortic arch (ARCH), descending thoracic aorta (DESC), infra-renal abdominal aorta (INFRARENAL), iliac artery (ILIAC), coronary artery (RCA), carotid artery (CAROTID), pulmonary artery (PA), and aortic root (ROOT)

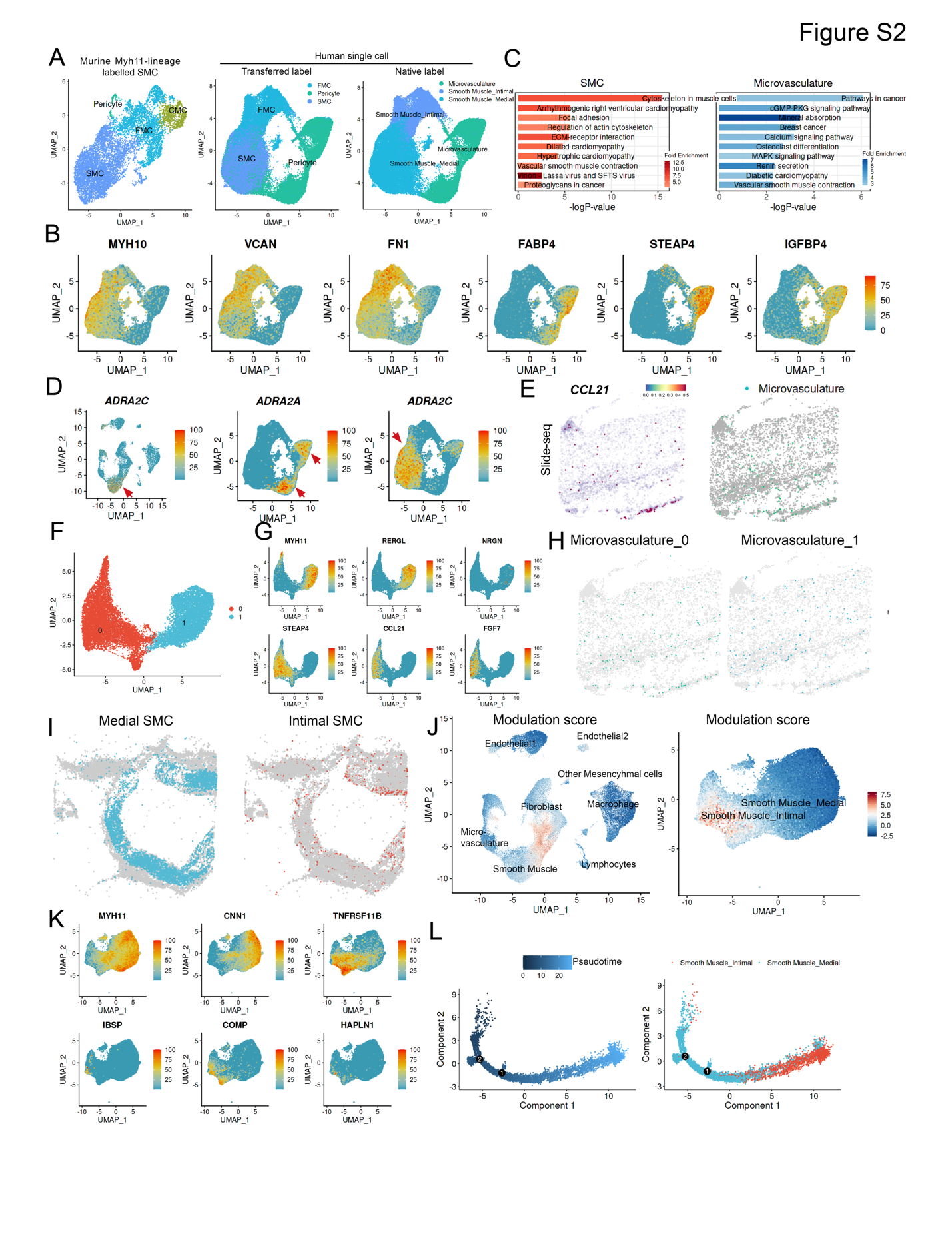
**Figure S2. Smooth muscle cell (SMC) lineage heterogeneity, and characterization of microvasculature and SMC subtypes.**

(A) UMAP plots demonstrating cross-species lineage consistency. Left: clustering of mouse smooth muscle cell (SMC) lineage cells. Middle: label transfer from mouse to human SMC lineage cells using supervised classification approaches. Right: native annotation of human SMCs based on canonical marker gene expression. The consistency among these three annotations supports conserved transcriptional programs across species.

(B) UMAP visualization of RNA expression levels of the differentially expressed genes in microvasculature (*FABP4, STEAP4 and IGBP4*) and smooth muscle cells (*MYH10, CAN and FN1*).

(C) Bar plot showing the z scores of enriched Kyoto Encyclopedia of Genes and Genomes (KEGG) pathways in differentially expressed genes between the two major SMC clusters (left: smooth muscle; right: microvasculature), indicating functional divergence in contractility, extracellular matrix remodeling, and signaling pathways.

(D) UMAP visualization of RNA expression levels of *ADRA2A* and *ADRA2C* in all cell clusters (left) or SMC-lineage clusters only (middle and right), highlighting their expression within subsets of the SMC population and suggesting subtype-specific adrenergic signaling regulation.

(E) Spatial distribution of *CCL21* expression in the aorta. Left: Slide-seq dim plot showing *CCL21* expression. Right: co-localization with microvasculature regions, indicating vessel-associated chemokine activity.

(F) UMAP showing the existence of two transcriptionally distinct subclusters within the microvasculature cell population.

(G) UMAP highlighting the top cluster-specific genes in the two microvasculature subclusters, reflecting functional specialization such as angiogenesis, permeability regulation, or immune interactions.

(H) Slide-seq spatial transcriptomics map illustrating that both microvasculature subclusters are predominantly localized to the adventitial layer of the aortic wall.

(I) Spatial Slide-seq plots depicting the anatomical localization of medial (left) and intimal (right) SMC populations in the right coronary artery (RCA), with distinct zonation patterns along the vessel wall.

(J) SMC modulation scores projected onto UMAPs of the full dataset (left) and SMC population only (right), highlighting the gradient of phenotypic modulation across SMCs.

(K) UMAP plots showing the RNA expression of classical contractile SMC markers (*MYH11*, *CNN1*) and markers associated with modulated SMC states (*TNFRSF11B*, *IBSP*, *HAPLN1*, *COMP*), underscoring the transcriptional plasticity of the SMC lineage.

(L) Single-cell trajectory analysis using Monocle reveals that the intimal (modulated) SMC population is positioned at a later stage along the pseudotime trajectory, originating from the mature (medial) SMC population.

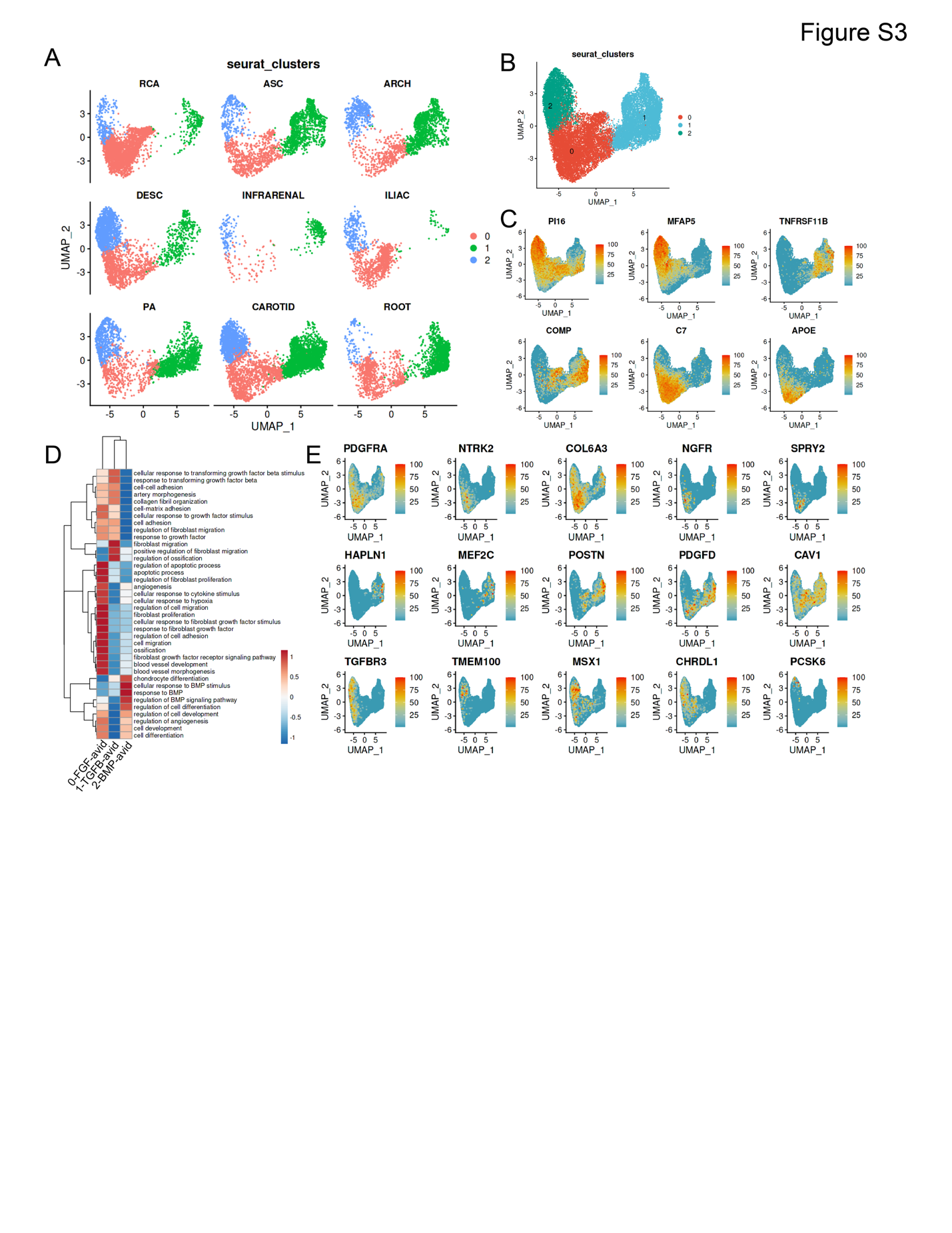

**Figure S3. Fibroblast heterogeneity, subclustering, and functional divergence across vascular segments.**

(A) Split-view UMAP plot displaying fibroblasts colored by segmental origin, revealing transcriptional heterogeneity associated with different vascular beds, such as ascending aorta, descending aorta, carotid, and coronary artery.

(B) UMAP embedding of the fibroblast cell population, showing the presence of three transcriptionally distinct subclusters.

(C) UMAP plot highlighting the expression of top cluster-specific genes within the three transcriptionally defined fibroblast subclusters, revealing distinct transcriptional programs associated with each subgroup.

(D) Heatmap showing the z scores of enriched functional pathways among differentially expressed genes across the three fibroblast subclusters. Enriched pathways include those related to ossification, TGF-β, and BMP signaling, indicating functional specialization among fibroblast subtypes.

(E) UMAP visualization of representative genes enriched in each fibroblast subcluster, highlighting key genes involved in TGF-β (top), ossification (middle), and BMP signaling (bottom).

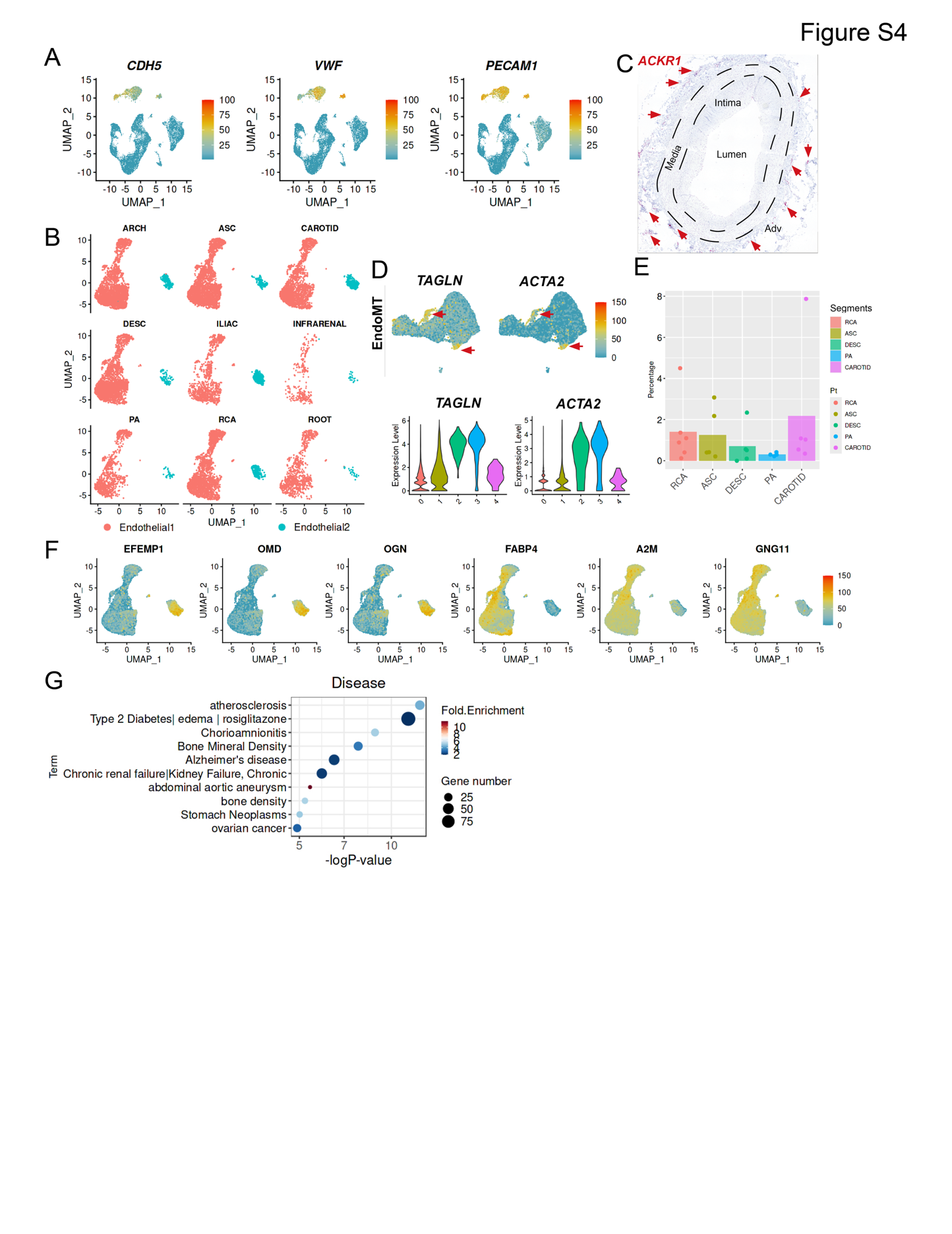

**Figure S4. Characterization of endothelial cell identity, regional heterogeneity, lineage features, and disease associations.**

(A) UMAP visualization showing the expression levels of canonical endothelial markers *CDH5*, *VWF*, and *PECAM1*, confirming the identity of the endothelial cell populations across the vascular beds.

(B) Split-view UMAP illustrating endothelial cells colored by their segmental origin, revealing transcriptional heterogeneity and regional specialization among vascular territories including aorta, coronary, and carotid arteries.

(C) RNAscope in situ hybridization detecting the expression of the cluster-specific gene *ACKR1* in the coronary artery, validating transcriptomic findings at the tissue level.

(D) UMAP plots (top) and corresponding violin plots (bottom) showing expression of non-endothelial lineage genes *TAGLN* and *ACTA2* within the Endothelial 1 population.

(E) Bar plot displaying the relative abundance of Endothelial 2 cells across four distinct vascular segments.

(F) UMAP visualization of RNA expression levels of the differentially expressed genes in Endothelial 1 (*EFEM1, OMD and OGN*) and Endothelial 2 (*FABP4, A2M and GNG11*).

(G) Dot plot showing the z scores of enriched GAD (Genetic Association Database) disease terms associated with the differentially expressed genes between Endothelial 1 and Endothelial 2 populations, indicating potential functional links between endothelial subtypes and human vascular pathologies.

Abbreviations: ascending thoracic aorta (ASC), aortic arch (ARCH), descending thoracic aorta (DESC), infra-renal abdominal aorta (INFRARENAL), iliac artery (ILIAC), coronary artery (RCA), carotid artery (CAROTID), pulmonary artery (PA), and aortic root (ROOT)

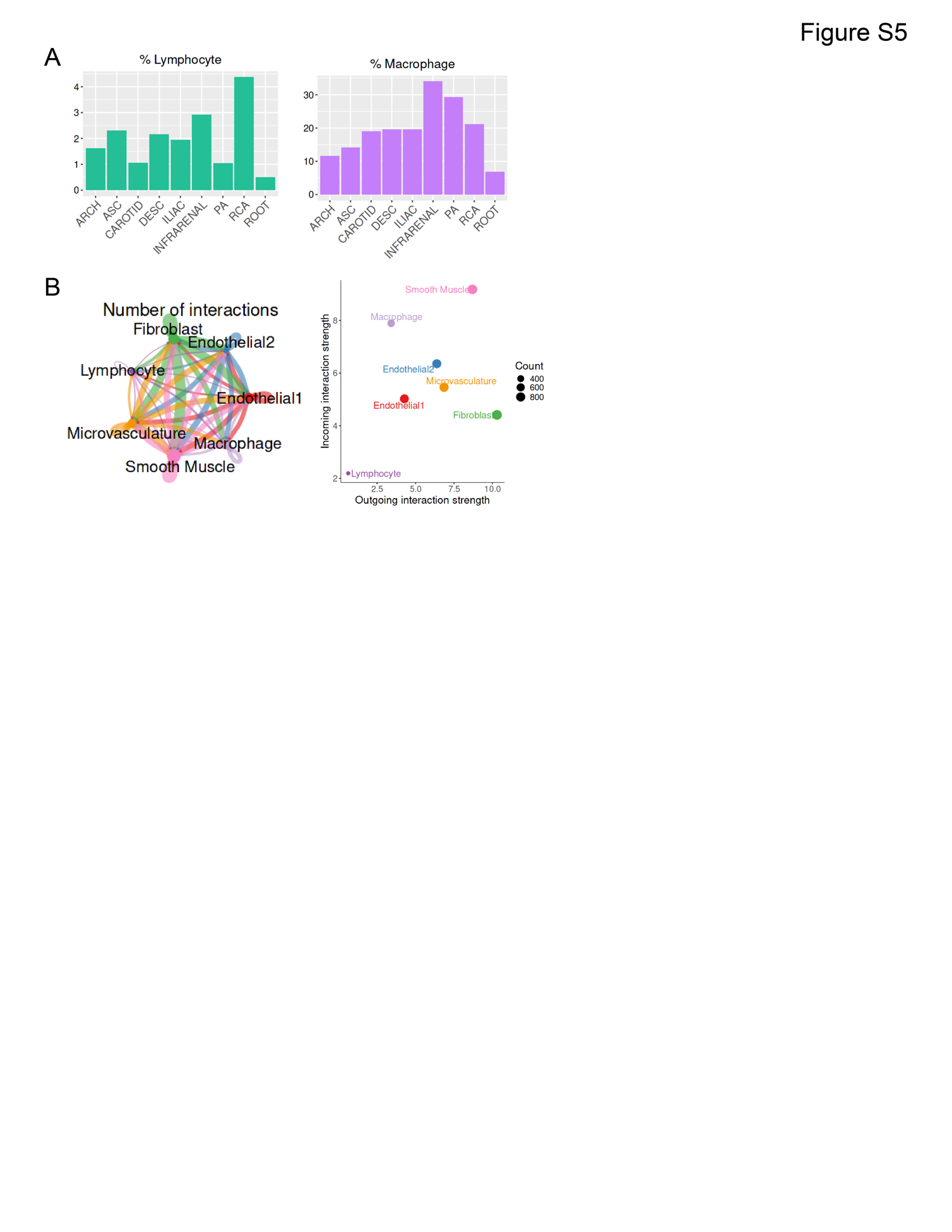

**Figure S5. Immune cell distribution and intercellular communication across human arterial segments.**

(A) Bar plots showing the fractions of lymphocytes (left) and macrophages (right) across distinct vascular segments, including ascending aorta, descending aorta, carotid artery, and coronary artery.

(B) Overview of predicted cell-cell communication networks among major arterial cell types inferred from ligand-receptor pair analysis. The left panel illustrates the global interaction network, where edge width represents the number of significant ligand-receptor interactions between each pair of cell types. The right panel presents a dot plot summarizing the directionality and strength of interactions, showing either incoming or outgoing signaling activity per cell type. Dot size reflects the number of interactions.

Abbreviations: ascending thoracic aorta (ASC), aortic arch (ARCH), descending thoracic aorta (DESC), infra-renal abdominal aorta (INFRARENAL), iliac artery (ILIAC), coronary artery (RCA), carotid artery (CAROTID), pulmonary artery (PA), and aortic root (ROOT)

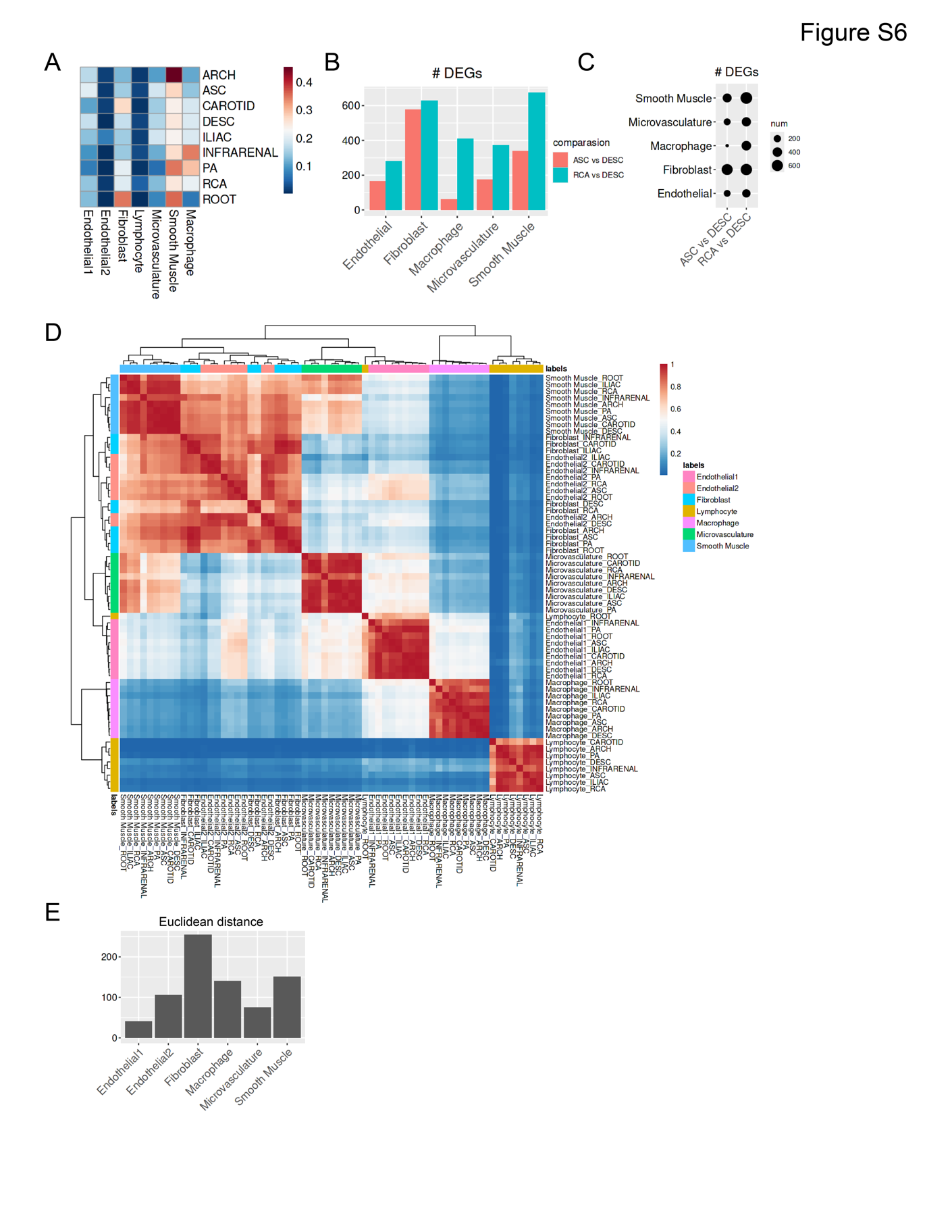

**Figure S6. Determinants of arterial identity across vascular territories.**

(A) Heatmap showing the relative proportions of distinct cell populations within each arterial bed, including ascending aorta, descending aorta, carotid artery, and coronary artery.

(B) Bar plot summarizing the number of differentially expressed genes (DEGs) identified between ascending vs. descending arteries and coronary vs. descending arteries across major vascular cell types. This highlights the extent of transcriptional divergence in each population.

(C) Dot plot displaying the DEG counts between ascending and descending arteries and between coronary and descending arteries across each cell type, with dot size indicating gene counts.

(D) Heatmap illustrating *Pearson* correlation coefficients among all arterial segments, based on global gene expression profiles aggregated across all cell types. Hierarchical clustering using *Euclidean* distance demonstrates transcriptomic relationships and anatomical lineage proximity between vascular regions.

(E) Bar plot showing the *Euclidean* distances between ascending and descending arterial sites for each cell type, providing a quantitative assessment of transcriptomic divergence that contributes to region-specific arterial identity.

Abbreviations: ascending thoracic aorta (ASC), aortic arch (ARCH), descending thoracic aorta (DESC), infra-renal abdominal aorta (INFRARENAL), iliac artery (ILIAC), coronary artery (RCA), carotid artery (CAROTID), pulmonary artery (PA), and aortic root (ROOT)

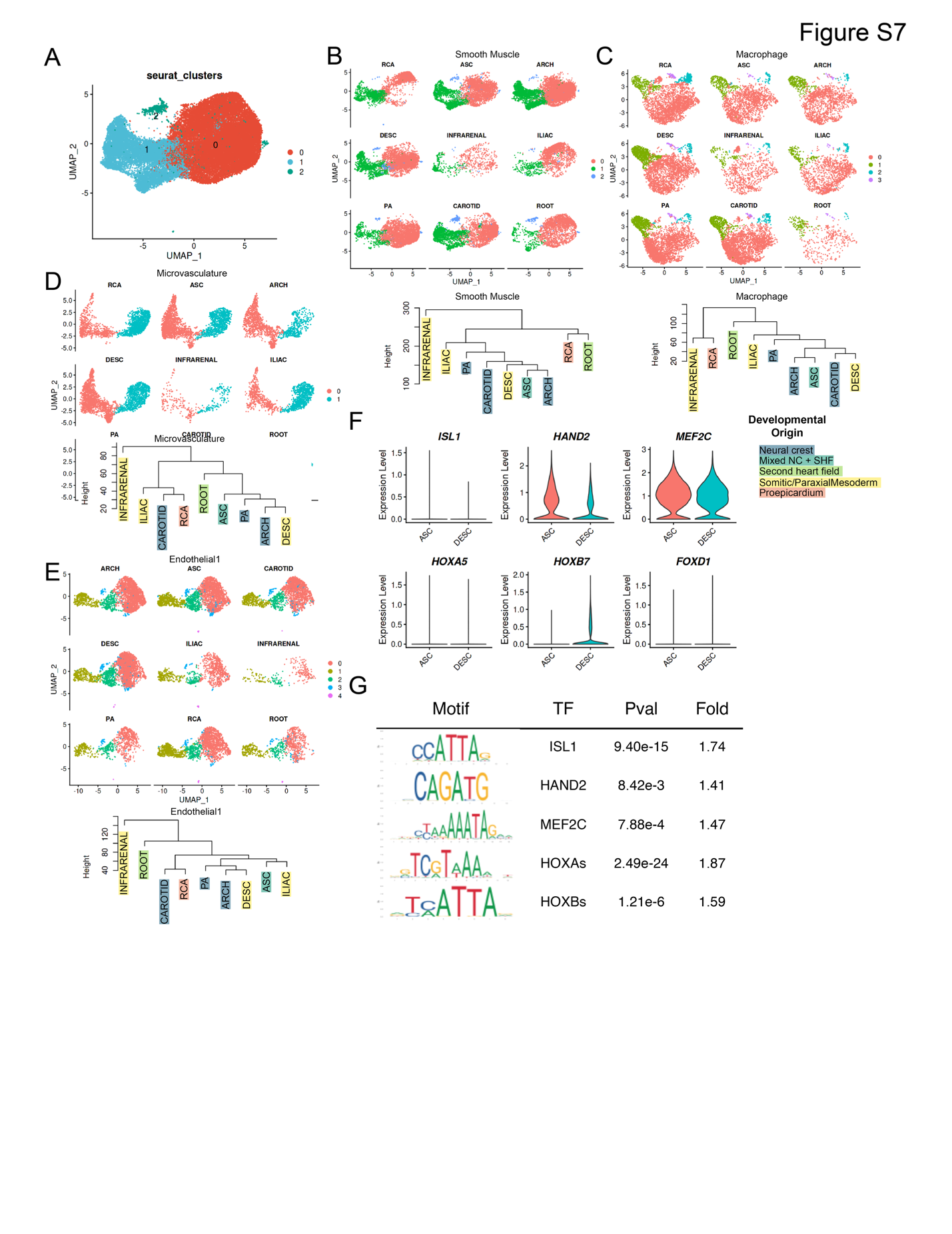

**Figure S7. The roles of embryonic origin in shaping transcriptomic diversity across arterial cell types.**

(A) UMAP embedding of smooth muscle cells (SMCs) reveals three transcriptionally distinct subclusters within the SMC lineage, indicating molecular heterogeneity.

(B) Top: Split-view UMAP showing the segmental origins of SMCs across the vasculature, demonstrating sgement-specific heterogeneity. Bottom: Dendrogram based on *Euclidean* distances calculated from the global transcriptome of SMCs, revealing clustering patterns that align with their anticipated embryonic lineage.

(C) Top: Segmental UMAP illustrating macrophages across arterial beds. Bottom: Dendrogram analysis of macrophage transcriptomes showing their correlation by segment.

(D) Top: Segmental UMAP of microvasculature cells. Bottom: *Euclidean* distance-based clustering of gene expression profiles shows concordance with embryonic origin in the microvasculature lineage.

(E) Top: Segment-specific UMAP visualization of endothelial1 cell. Bottom: Dendrogram analysis of endothelial1 transcriptomes, ordered by embryonic lineage correlation across arterial regions.

(F) Violin plots showing segment-specific expression of transcription factors associated with embryonic patterning, including *ISL1*, *HAND2*, *MEF2C*, and representative *HOX* genes in SMCs derived from ascending and descending arteries.

(G) Motif enrichment analysis showing significant overrepresentation of ISLET1, HAND2, MEF2C, and HOX transcription factor (TF) binding sites in the promoter regions of differentially expressed genes between ascending and descending SMCs. Macrophages serve as a negative control to underscore lineage-specific regulatory enrichment in SMCs.

Abbreviations: ascending thoracic aorta (ASC), aortic arch (ARCH), descending thoracic aorta (DESC), infra-renal abdominal aorta (INFRARENAL), iliac artery (ILIAC), coronary artery (RCA), carotid artery (CAROTID), pulmonary artery (PA), and aortic root (ROOT), Neural crest (NC), Second heart field (SHF)

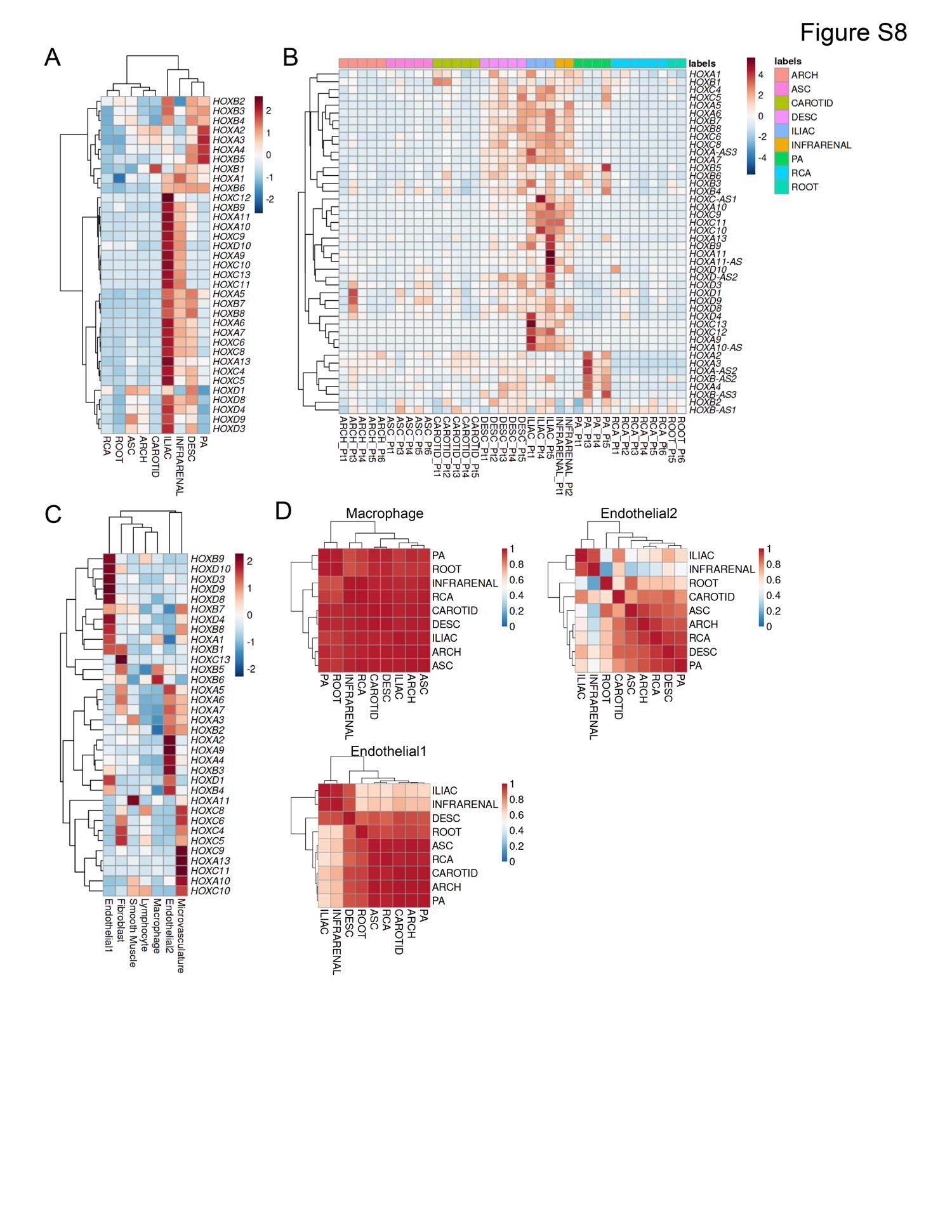

**Figure S8. Segment and cell type-specific patterns of vascular HOX gene expression.**

(A) Heatmap depicting the standardized expression (z scores) of *HOX* genes across distinct vascular segments, revealing transcriptional signatures along the arterial axis.

(B) Patient-specific heatmap illustrating the z-scored expression of *HOX* genes across various cell types in six individuals (Pt1–Pt6).

(C) Heatmap showing the z scores of *HOX* gene expression across all major cell types, emphasizing lineage-dependent transcriptional programs linked to positional identity.

(D) Heatmaps of *Pearson* correlation coefficients among arterial segments based on *HOX* gene expression profiles, with hierarchical clustering performed using *Euclidean* distance. The analysis is shown for macrophages (top left), endothelial1 cells (bottom left), and endothelial2 cells (top right).

Abbreviations: ascending thoracic aorta (ASC), aortic arch (ARCH), descending thoracic aorta (DESC), infra-renal abdominal aorta (INFRARENAL), iliac artery (ILIAC), coronary artery (RCA), carotid artery (CAROTID), pulmonary artery (PA), and aortic root (ROOT)

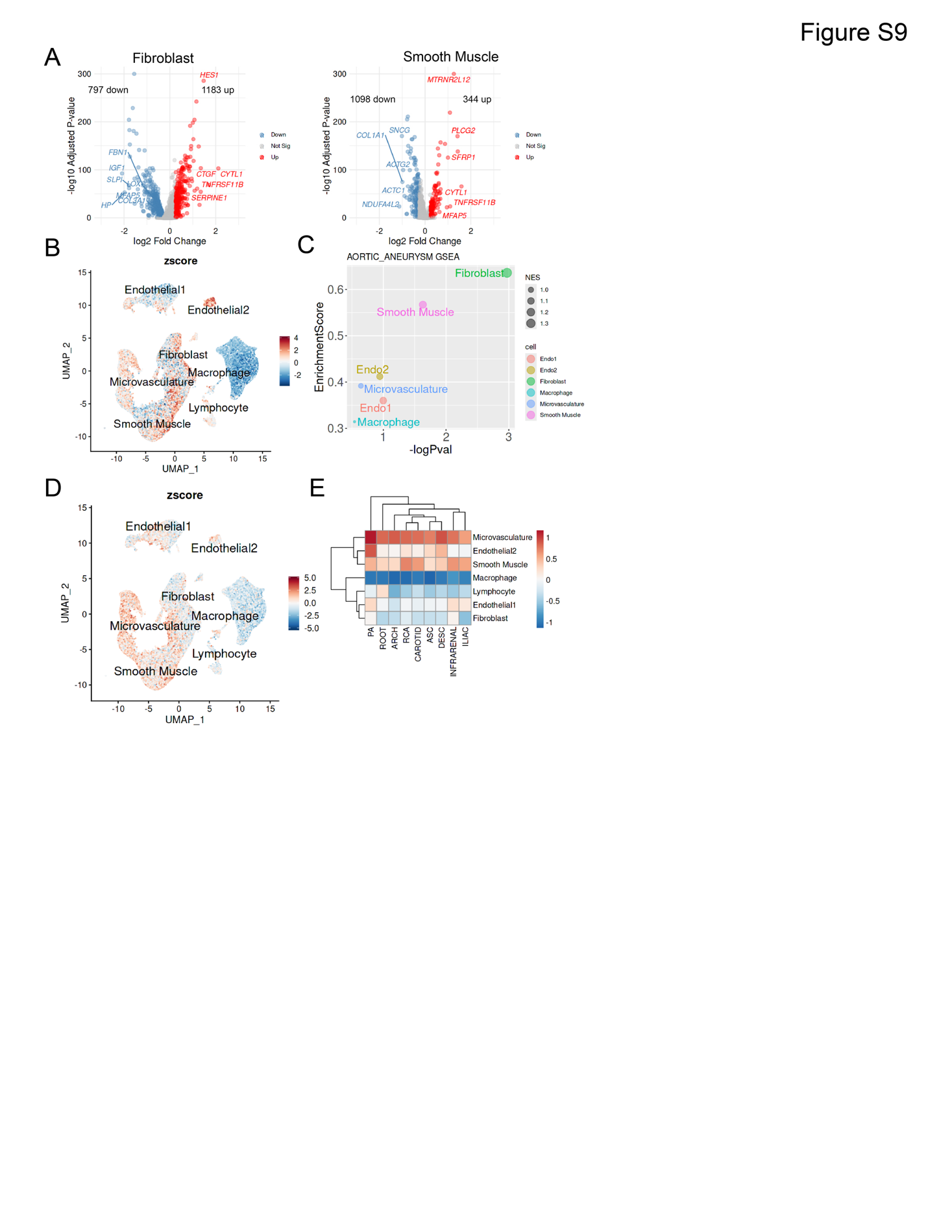

**Figure S9. Cell type-specific enrichment of aortic aneurysm-associated and GWAS signals.**

(A) Differentially expressed genes (DEGs) between the descending and ascending aortic fibroblast cell (left) or smooth muscle cell (right), showing average log₂ fold change and -log10(P-values), indicating transcriptional divergence and potential functional specialization.

(B) UMAP visualization of single-cell disease-relevance score (scRDS), mapping the inferred disease relevance of individual cells based on RNA expression profiles and aortic aneurysm GWAS genes.

(C) Gene Set Enrichment Analysis (GSEA) of genome-wide association study (GWAS) genes linked to aortic aneurysm, performed against differentially expressed genes from all major vascular cell types. The analysis highlights significant enrichment in smooth muscle cells (SMCs) and fibroblasts, suggesting their predominant contribution to disease susceptibility.

(D) UMAP visualization of scRDS, mapping the inferred disease relevance of individual cells based on RNA expression profiles and Million Veteran Program (MVP) GWAS genes.

(E) Heatmap showing the relative scRDS scores of D for each cell type across various vascular segments.

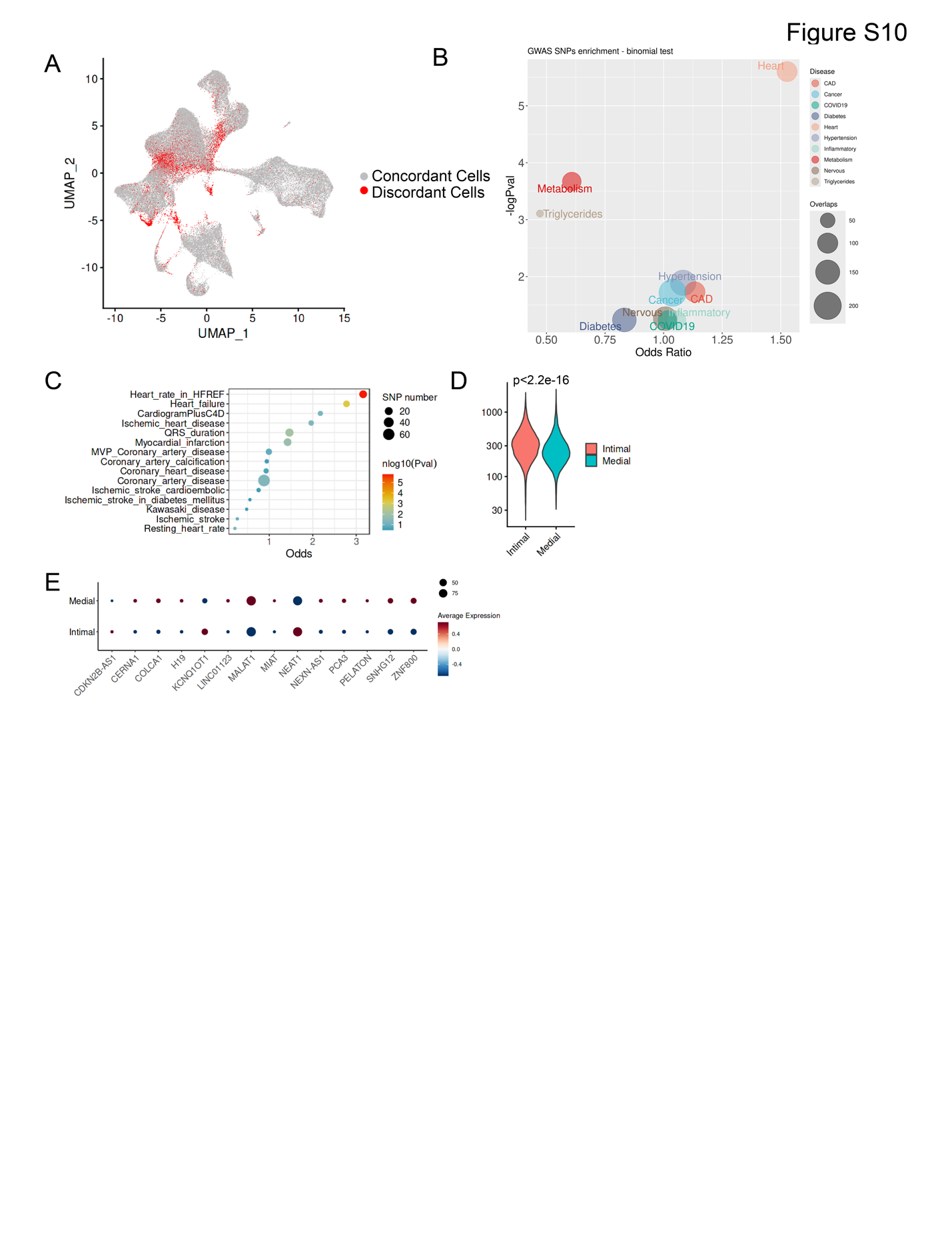

**Figure S10. lncRNA expression patterns and their genetic associations with vascular disease.**

(A) UMAP visualization and clustering of single cells based on the sc-lncRNA transcriptome, annotated using both whole-transcriptome-derived and lncRNA-derived cell type identities. Highlighted regions indicate discordant annotations.

(B) Bubble plot showing the enrichment of overlap between differentially expressed lncRNAs and GWAS catalog SNPs across a range of diseases. Notably, vascular-related diseases such as coronary artery disease (CAD) show strong associations.

(C) Dot plot representing the binomial enrichment of differentially expressed lncRNAs across GWAS disease categories. Terms related to cardiac and coronary conditions are significantly enriched, indicating potential regulatory roles of lncRNAs in vascular pathogenesis.

(D) Violin plot comparing the number of expressed lncRNA genes (TPM > 0) in intimal versus medial SMC populations. The results demonstrate increased lncRNA transcriptional activity in the modulated (intimal) SMCs, consistent with their disease-relevant phenotypic state.

(E) Dot plot showing the expression levels of previously reported CAD-associated lncRNAs, differentially expressed between intimal and medial SMCs. These findings provide functional context for lncRNAs implicated in vascular disease through human genetics.

Abbreviations: genome wide association study (GWAS), Coronary artery diseases (CAD), heart failure with reduced ejection fraction (HFREF)
